# An embodied perspective: angular gyrus and precuneus decode selfhood in memories of naturalistic events

**DOI:** 10.1101/2024.09.09.612088

**Authors:** Heather M. Iriye, Peggy L. St. Jacques

## Abstract

While we often assume that memory encoding occurs from an in-body (first-person) perspective, out-of-body experiences demonstrate that we can form memories from a third-person perspective. This phenomenon provides a distinctive opportunity to examine the interaction between embodiment and visual perspective during encoding, and how this interplay shapes the recall of past events. Participants formed memories for naturalistic events following a manipulation of their sense of embodiment from in-body and out-of-body perspectives and recalled them during functional scanning. Region of interest multivariate analyses examined how the angular gyrus, precuneus, and hippocampus reflected visual perspective, embodiment, and their interaction during remembering. Patterns of activity during retrieval in the left angular gyrus and bilateral precuneus predicted embodiment on its own separated from visual perspective. In contrast, we observed only inconclusive evidence that these posterior parietal regions predicted visual perspective independent of embodiment. While the left angular gyrus distinguished between in-body and out-of-body perspectives during the retrieval of events associated with both strong and weak embodiment, decoding accuracy predicting visual perspective was only above chance for events encoded with strong embodiment in the precuneus bilaterally. Our results suggest that the contribution of posterior parietal regions in establishing visual perspectives within memories is tightly interconnected with embodiment. Encoding events from an embodied in-body perspective compared to embodied out-of-body perspective led to higher memory accuracy following repeated retrieval. These results elucidate how fundamental feelings of being located in and experiencing the world from our own body’s perspective are integrated within memory.

---

The ability to re-experience past events vividly and accurately through remembering is tightly linked to the awareness and perception of one’s body, or bodily selfhood (e.g., (Bergouignan, Nyberg, & Ehrsson, 2014; Bréchet et al., 2020, 2019; Iriye & Ehrsson, 2022; Penaud, Yeh, Gaston-Bellegarde, & Piolino, 2023). Bodily selfhood typically involves the feeling of being located in one’s body at a specific point in space (i.e., self-location), a sense of ownership over one’s body (i.e., body ownership), and experiencing the world from an in-body, first-person perspective (Blanke et al., 2015). However, disturbances to the sense of bodily selfhood such as observer experiences, or out-of-body experiences, arise whereby one senses a detachment from the physical body during an event and is afforded a view of “how the entire scene would appear … to an onlooker who sees us as well as our surroundings” (p. 469, Nigro & Neisser, 1983). Observer and out-of-body experiences are often reported in the contexts of schizophrenia and post-traumatic stress disorder (Blackmore, 2019; Reynolds & Brewin, 1999). Yet, 10-15% of the general population is estimated to have at least one observer experience over their lifetime (Blackmore, 2017, Blanke et al., 2004), leading to them to be described as “normal though unusual experiences” (p. 163, (Amorim, 2003). Highly emotional situations that involve a strong degree of self-awareness may lead to observer/out-of-body experiences (Nigro & Neisser, 1983). For example, while giving a high-pressure presentation to a large crowd, an individual may spontaneously see themselves as they appear to an audience member sitting in the audience. It remains unknown how alteration in the sense of self-location, body ownership, and visual perspective during encoding affect neural mechanisms of memory retrieval. For example, most neuroimaging studies have focused on manipulations of visual perspective during memory retrieval (Eich et al., 2009a; Freton et al., 2014a; Grol et al., 2017a; Hebscher et al., 2021; St. Jacques et al., 2017, 2018) rather than on encoding (but see Bergouignan et al., 2014). Here, we employ an innovative paradigm that allows the manipulation of visual perspective (i.e., whether an event is formed from either a natural in-body perspective or an out-of-body perspective) alongside feelings of self-location and body ownership (i.e., embodiment) during the formation of highly realistic events to investigate how these essential aspects of selfhood manifest in patterns of neural activity at retrieval. This research develops a new comprehension of how basic forms of selfhood rooted in the body structure higher-level aspects of selfhood tightly linked with episodic memory (Iriye & St. Jacques, 2019; Prebble et al., 2013). Clinically, insight into the relationship between visual perspective, embodiment and memory has the potential to deepen understanding of why disorders characterized by abnormal body experiences like dissociation from one’s physical body, such as is the case in schizophrenia, also incur episodic memory deficits (Borda & Sass, 2015; Prebble et al., 2013; Sierra & David, 2011).

The sense of bodily selfhood is experimentally manipulated by leveraging the brain’s multisensory mechanisms that underlie own-body perception (e.g. Guterstam, Björnsdotter, Gentile, et al., 2015a; Petkova & Ehrsson, 2008; van der Hoort et al., 2011). In these experiments, participants wear a virtual reality head-mounted display (HMD) unit connected to a camera attached to a mannequin, which allows the participant to view the lab from the mannequin’s perspective. Then the experimenter applies brushstrokes to both the mannequin and correlated locations on the participant’s real body. When brushstrokes are synchronous (i.e., seen and felt touches occur at the same time and in the same location), the congruence of visual and tactile information induces a sense of embodiment over the mannequin. Conversely, asynchronous brushstrokes do not lead to illusory embodiment over the mannequin due to the mismatch between visual and tactile information. Questionnaires and physiological recordings (e.g., skin conductance responses) measure the strength of these bodily illusions (e.g. Guterstam & Ehrsson, 2012; O’Kane & Henrik Ehrsson, 2021). This paradigm has been extended to create a sense of ownership over an invisible body, highlighting the exceptionally flexible nature of bodily selfhood (D’Angelo et al., 2017; Guterstam, Abdulkarim, et al., 2015; Kondo et al., 2018). For example, Guterstam and colleagues (2015) and D’Angelo and colleagues (2017) had participants wear HMD units mounted to a wall, allowing them to see empty space below the cameras. The experimenter induced a sense of embodiment over an invisible body by applying synchronous brushstrokes to the participant’s limbs and torso and the corresponding location in empty space below the camera. Asynchronous visuotactile stimulation did not induce the invisible body illusion. However, participants’ real bodies were not visible in the scene in these studies. In contrast, in the current study we implemented the invisible body illusion to manipulate bodily selfhood from in-body and out-of-body perspectives that incorporated participants’ own physical bodies in their visual field, replicating clinical reports of out-of-body experiences defined by seeing one’s body from a third-person perspective (Blanke et al., 2004).

Previous research suggests that the individual components of bodily selfhood (i.e., body ownership, self-location, and visual perspective) may be supported by activity within the angular gyrus, precuneus, and hippocampus. Iriye, Chancel, and Ehrsson (2024) immersed participants in naturalistic scenes involving lifelike events while manipulating ownership over a mannequin’s body aligned with participants’ real bodies with synchronous and asynchronous visuotactile stimulation during functional magnetic resonance imaging (fMRI) scanning. One week later, participants retrieved these events from memory also during fMRI scanning. The authors observed that patterns of activity in regions including the left angular gyrus, right precuneus, and bilateral hippocampi predicted whether body ownership was strong (i.e., synchronous visuotactile stimulation) compared to weak (i.e., asynchronous visuotactile stimulation) during encoding. Moreover, memory reinstatement in these three regions, as measured by encoding-retrieval pattern similarity, was greater for lifelike memories formed with strong body ownership and high levels of memory vividness, compared to weak body ownership and low memory vividness. Other research has indicated that a portion of the left angular gyrus, linked to multisensory integration and visuospatial representations of body parts (Vingerhoets, 2014), is implicated in implementing probabilistic models that dynamically and continuously generate the sense of body ownership by comparing incoming sensory information with previous perceptual experiences (Chancel et al., 2022). Interfering with activity in the angular gyrus through brain lesions (Ionta et al., 2011), seizures (Blanke et al., 2004), or transcranial magnetic stimulation (Blanke et al., 2002) is linked to out-of-body experiences involving the feeling of dissociation from one’s physical body and a shift in visual perspective to an observer/out-of-body point of view. Additionally, damage to the angular gyrus is linked to decreased use of in-body perspective imagery while mentally navigating (Ciaramelli et al., 2010a). Consistent with neuropsychological findings, continuous theta burst stimulation to the left angular gyrus decreases the likelihood of recalling memories from an in-body perspective (Bonnici et al., 2018a). However, the angular gyrus may play a role in establishing both in-body and out-of-body perspectives. St. Jacques, Szupunar, and Schacter (2017) observed that alternating between visual perspectives is linked to increased activity in the right angular gyrus, indicating its role in representing both in-body and out-of-body perspectives. Additionally, Grol, De Raedt, and Vingerhoets (2017) observed heightened activity in this region when recalling memories from an out-of-body versus in-body perspective, likely linked to mental transformations necessary for updating visual perspective. Therefore, additional research is necessary to clarify the role of the angular gyrus in establishing egocentric perspectives during memory retrieval. Current evidence suggests that multisensory information underpinning one’s sense of bodily selfhood is combined with additional multi-modal memory components within a unified egocentric perspective in the angular gyrus.

While egocentric perspectives are established in the angular gyrus, they are manipulated in the precuneus (St. Jacques et al., 2017, 2018). The precuneus is linked to visual mental imagery (Cavanna & Trimble, 2006; Fletcher et al., 1995) and the transformation of allocentric spatial representations stored in the hippocampus into egocentric reference frames (Byrne et al., 2007). Several studies have reported that adopting an out-of-body perspective during memory retrieval recruits the precuneus (Faul et al., 2020; Grol et al., 2017; St. Jacques et al., 2017). In contrast, structural neuroimaging studies have reported a positive correlation between gray matter volume in the precuneus and the tendency to use in-body perspective imagery when retrieving memories (Grol et al., 2017; Hebscher et al., 2018). These conflicting findings may be explained by considering that the precuneus contributes to the construction of mental scenarios from an egocentric framework, such that adopting a more atypical observer perspective places greater demands on the involvement of this region (Iriye & St. Jacques, 2020). According to this interpretation (St. Jacques, 2019), the precuneus facilitates both own eyes and observer perspectives during memory retrieval. Supporting this idea, St. Jacques and colleagues (2018) reported common involvement of the precuneus when adopting a novel observer and own eyes perspective. Nonetheless, greater research is needed to elucidate how different visual perspectives are represented in this region. In addition to visual perspective, the precuneus may also play a key role in bodily selfhood more widely. For example, direct cortical stimulation of the anterior precuneus leads to altered bodily sensations such as falling/dropping, floating, dizziness, and self-dissociation (Lyu et al., 2023). Together these findings imply that the precuneus is involved in the ability to construct mental scenes from an embodied perspective during memory retrieval.

Beyond the parietal cortex, the hippocampus is implicated in both embodiment and episodic memory. Scenes encoded with a preserved sense of agency, a key facet of embodiment, are reinstated during retrieval to a greater degree in the hippocampus, which predicts recognition memory accuracy and is coupled with reinstatement effects in the premotor cortex (Meyer et al., 2024). Further, the hippocampus has been shown to contain information about the sense of self-location (Guterstam, Björnsdotter, Bergouignan, et al., 2015; Guterstam, Björnsdotter, Gentile, et al., 2015b), which may affect memory encoding and retrieval.(Meyer et al., 2024). Consistent with this idea, Bergouignan and colleagues (2014) reported repetition enhancement effects in the left posterior hippocampus during repeated retrieval of memories formed from an out-of-body perspective. The strength of this hippocampal repetition enhancement effect predicted concomitant reductions in reported memory vividness (i.e., clarity of mental images during remembering) and impaired recall of episodic details for memories encoded from an out-of-body compared to in-body perspective. Conversely, repeated retrieval of realistic memories encoded from a more natural in-body perspective was linked to repetition suppression effects in the left posterior hippocampus. The authors suggested that encoding memories from out-of-body perspectives interrupts hippocampal encoding mechanisms, which affects activation patterns during repeated memory retrieval. However, participants in this experiment experienced only embodied in-body and out-of-body perspectives, induced by synchronous visuotactile stimulation, and thus, it is unclear whether the hippocampus would be sensitive to feelings of disembodiment during memory formation, induced by asynchronous visuotactile stimulation. In the present study we tested whether the hippocampus contained separate neural representations of embodied viewpoints using a two (embodiment: synchronous, asynchronous) x two (visual perspective: in-body, out-of-body) experimental design and a multivariate analytical approach, which is thought to reflect neural representations in a similar way to repetition suppression (Barron et al., 2016). Collectively, earlier research indicates that the hippocampus might be responsive to the combined effects of visual perspective, self-location, and body ownership when recalling events encoded from in-body and out-of-body perspectives with the aim of supporting spatial memory processes.

We conducted the present study to investigate how visual perspective (i.e., whether an event is experienced from an in-body or out-of-body perspective) and the sense of embodiment (i.e., here body ownership and self-location) interact to structure patterns of activity in the angular gyrus, precuneus, and hippocampus during memory retrieval. Participants encoded memories for a set of realistic events following an established paradigm to manipulate their sense of bodily self from in-body and out-of-body perspectives using synchronous and asynchronous visuo-tactile stimulation. There were four experimental conditions: in-body synchronous, in-body asynchronous, out-of-body synchronous, out-of-body asynchronous. Memories for these events were repeatedly retrieved during functional magnetic resonance imaging (fMRI) scanning later that same day. We trained multivariate classifiers to predict whether patterns of activity within the three targeted regions of interest reflected visual perspective (i.e., in-body versus out-of-body perspectives, collapsed across embodiment conditions), embodiment (i.e., synchronous versus asynchronous visuotactile stimulation, collapsed across visual perspectives), and their interaction (i.e., synchronous in-body versus synchronous out-of-body and asynchronous in-body versus asynchronous out-of-body) during retrieval.

Regarding the angular gyrus, we hypothesized that patterns of activity would reflect visual perspective (i.e., in-body, out-of-body), sense of embodiment (i.e. synchronous, asynchronous), and their interaction based on this region’s involvement in multisensory integration of bodily signals underpinning own-body perception (i.e., Iriye et al., 2024; Chancel et al., 2023) and participation in establishing both in-body and out-of-body egocentric perspectives (Bonnici et al., 2018; Grol et al., 2017; St. Jacques et al., 2017). Concerning the precuneus, we expected that patterns of activity during memory retrieval would reflect the sense of embodiment present during encoding of the event, due to recent research linking stimulation of this region to altered states of bodily awareness (Lyu et al., 2023). We also predicted that the activity in the precuneus would not represent visual perspective on its own separated from embodiment, as previous research has shown that the precuneus supports retrieval from both in-body and out-of-body perspectives (e.g., St. Jacques et al., 2018). Instead, we expected that the patterns of activity in the precuneus would reflect the interaction between visual perspective and embodiment as direct stimulation of this region leads to dissociative bodily states (Lyu et al., 2023), which are more commonly linked to out-of-body compared to in-body perspectives (Blanke et al., 2004). For the hippocampus, we predicted that patterns of activity in this region would reflect visual perspective, embodiment, and their interaction due to reports of its involvement in memory studies manipulating embodiment (Iriye et al., 2024), and visual perspective and embodiment together (Bergouignan et al., 2014).

## Methods

### Participants

Participants included 28 healthy, right-handed young adults (age range: 18 to 29 years), with no prior history of neurological or psychiatric impairment, and who were not currently taking medication that affected mood or cognitive function. Participants provided informed written consent, approved by the School of Psychology at the University of Sussex, prior to their participation in the study. Functional scanning data from four participants was excluded due to a technical issue. Thus, the final analysis was performed on 24 participants (14 women, 10 men; *mean age* = 20.46, *SD* = 2.21; *mean years of education* = 14.96, *SD* = 1.68). The sample size in the present study (*N* = 24) is comparable with the sample size of previous neuroimaging studies investigating the role of bodily selfhood on memory (e.g., Bergouignan et al., 2014: *N* = 21; Iriye et al., 2024: *N* = 24). Ethical approval was obtained by the Sciences & Technology Cross-Schools Research Ethics Committee (SCITEC C-REC) at the University of Sussex.

### Procedure

#### Overview

The experiment involved five main phases: illusion induction, event encoding, immediate memory test, retrieval during fMRI scanning, and a post-scanning memory test. The first part of the experiment involved altering participants’ sense of bodily ownership, self-location, and embodiment with two well-established illusion inductions known as the full-body illusion (e.g., Tacikowski, Weijs, & Ehrsson, 2020) and a variant of the invisible body illusion (e.g., Guterstam, Abdulkarim, et al., 2015). Participants wore a virtual reality head mounted display (HMD) that received input from a camera mounted to either a tripod positioned behind the participant at eye level, such that they could view their physical body as if looking at themselves from behind (i.e., out-of-body perspective), or to the participant’s head such that they viewed their physical body from a first-person vantage point when they looked down (i.e., in-body perspective). Synchronous visuotactile stimulation was used to induce a feeling of ownership over the participant’s real body in the in-body perspective, or an invisible body located in empty space below the cameras in the out-of-body perspective. Asynchronous visuotactile stimulation reduced/abolished ownership over the participant’s real body (in-body perspective) or the invisible body located beneath the cameras (out-of-body perspective). Thus, the experiment involved a 2 (visual perspective: in-body, out-of-body) x 2 (embodiment: synchronous, asynchronous) within-subjects design (see Figure 1A&B). After the first illusion had been induced, the event encoding task involved asking participants to engage in a series of short, lifelike events comprised of four separate word games played with the experimenter. Then, during an immediate memory test, participants practiced vividly retrieving each game with their closed eyes immediately after each game was complete. They also answered questions relating to (1) the strength of the illusion induction, (2) details of each event as a measure of memory accuracy, and (3) subjective ratings of own eyes perspective taking, observer perspective taking, vividness, emotional intensity, and perceived memory accuracy on seven-point Likert scales. This procedure was then carried out for the other three conditions. Later that day, participants retrieved memories for the word games during fMRI scanning. They were asked to repeatedly retrieve memories for each event/word game, and rate visual perspective (on separate in-body and out-of-body scales) and vividness. Finally in a post-scanning memory test, they were asked additional novel questions about each event and completed ratings of emotional intensity and perceived memory accuracy. Below we discuss each of the experimental phases in detail.

**Figure 1.**
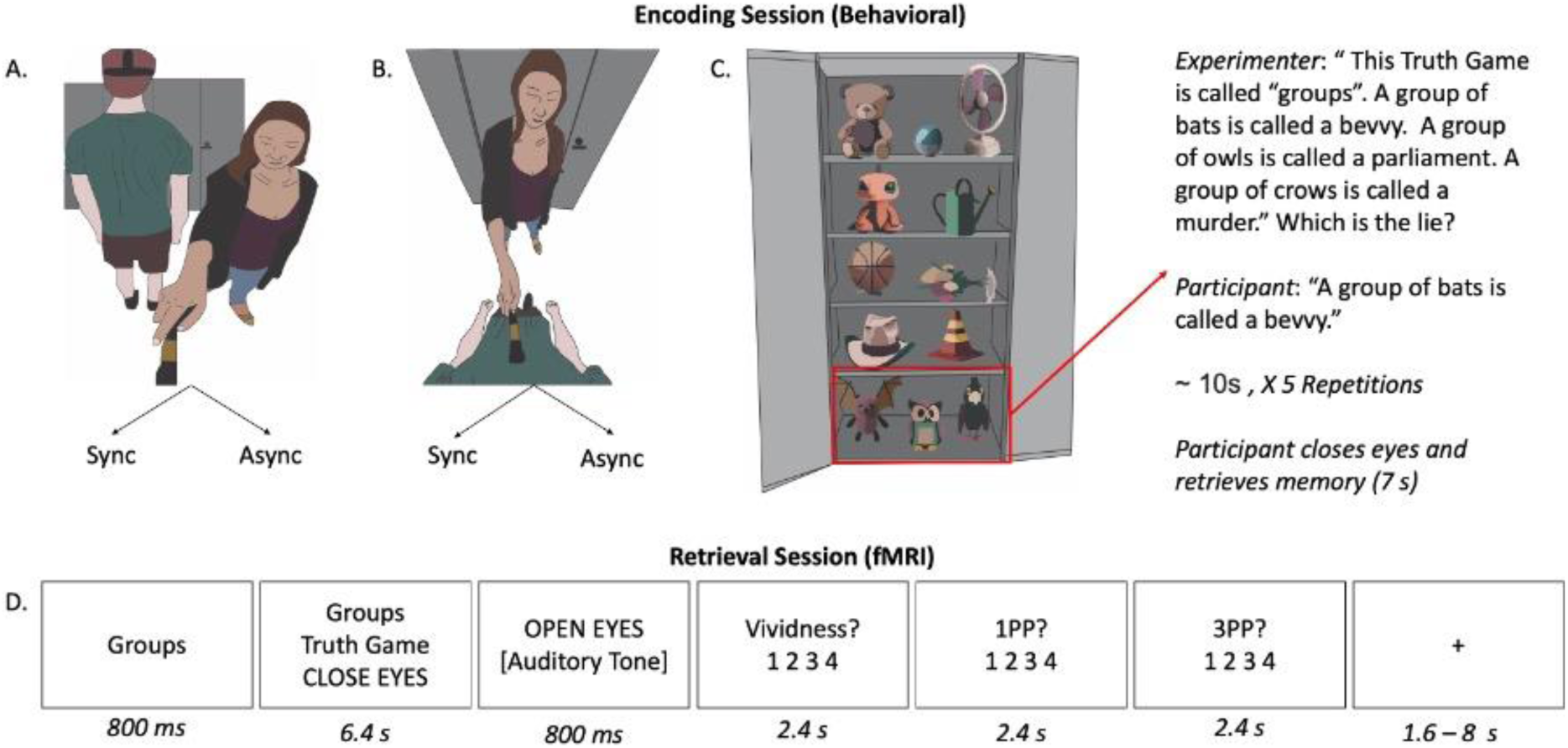
Participants wore a virtual HMD and viewed a live feed of the experimental room from either an out-of-body. (A) or in-body perspective (B). In half of the trials, participants received synchronous (sync) visuotactile stimulation to induce a sense of ownership either over their own body in the own eyes condition, or an invisible body located under the camera in the observer condition. In the other half of the trials, the experimenter administered asynchronous (async) visuotactile stimulation to reduce the sense of ownership over the participant’s real body (in-body perspective) or the invisible body located below the camera (out-of-body perspective). The participants then played a series of word games with the experimenter based on unique items located in the cabinet in front of them (C). There were four games in each round (Truth Game, I Spy, Categories, Sentence Construction) and each game was repeated a total of five times. At the end of the last repetition of each game, participants closed their eyes and retrieved a memory for the game for 7 s before moving on to the next game. At the end of each round of four games, participants completed a cued recall accuracy test for each word game and subjective ratings of vividness, emotional intensity, and the degree to which they retrieved their memory from a 1PP and 3PP on separate scales. Later that day, participants retrieved memories for the word games during functional scanning and rated the vividness of their mental imagery and the perspective adopted during retrieval on separate own eyes and observer scales (D). Participants completed a second cued recall test and rated the emotional intensity and belief in memory accuracy associated with their memories post-scanning.

#### Illusion Inductions

Two videos were recorded for the illusion induction, one from an out-of-body perspective and one from an in-body perspective. For the out-of-body perspective video, a high definition 360-degree camera (i.e., Ricoh Theta S; Resolution: 1920 x 1080; Frame Rate: 29.97 frames per second) was mounted on a tripod, adjusted to the participant’s eye-level, and placed behind the participant. Next, participants were positioned to face a set of closed cabinets on an “X” marked on the floor in the lab and asked to wear an Oculus Rift HMD. A tripod was placed one meter behind the participant. The HMD display remained blank during video recording. The position of the camera on the tripod created an out-of-body perspective by presenting a view of the participant’s back body, as if they were standing behind themself. After video recording started, the experimenter approached the tripod and stroked empty space below the camera with a medium sized paintbrush in locations corresponding to the participant’s torso, arms, and legs, as if applying brushstrokes to an invisible body positioned below the camera (Guterstam, Abdulkarim, et al., 2015). The length of the participant’s torso, arms, and legs were marked against a wall located behind the tripod to indicate starting and stopping points for each brushstroke, ensuring that brushstrokes matched the participant’s specific body dimensions. Each brushstroke lasted one second with an additional one and a half seconds between brushstrokes (Guterstam, et al., 2015). Five brushstrokes were applied to each of the five different body parts in the following order: torso, right arm, left arm, right leg, left leg. After the first video had been recorded, the camera was mounted to a small, flexible tripod attached to the front of the HMD unit to create a video from an in-body perspective (see Figure 1B). Then, the experimenter applied brush strokes to the participant’s physical body using the same procedure as the first video. After both videos had been recorded, they were converted to a 360-degree MP4 format using the Ricoh Theta desktop application while the participant waited outside the lab. Videos were presented using Whirligig software (Free Version; Fleischman, 2000), which allowed them to be visible both inside the HMD unit and on the desktop computer screen. Each video was 62.5 seconds in length.

Participants were then invited back into the laboratory where they were once again fitted with the HMD unit and stood on the “X” marked on the floor facing the cabinets. Participants were instructed to look down at their body. Masking tape was placed on the underside of the HMD unit to ensure that the participants only saw what was presented on the screen without any additional light entering the display. The experimenter then played the video visible to the participant through the HMD unit and the experimenter through the desktop computer to the right of the participant. The experimenter applied brushstrokes to the participant’s body either synchronously or asynchronously with the timing of brushstrokes in the video in the same location as seen in the video. Synchronous (i.e., sync) seen and felt brushstrokes induce a sense of ownership over the participant’s physical body for the in-body perspective (Tacikowski et al., 2020) and what is perceived to be an invisible body located below the cameras in the out-of-body perspective (D’Angelo, Di Pellegrino, & Frassinetti, 2017; Guterstam et al., 2015; Kondo et al., 2018). Note that the key difference between the typical invisible body illusion and the present study design is that the participant’s physical body is seen from behind in the current experiment, affording an out-of-body perspective on the visual scene. Previous research has shown that individuals retain a strong sense of self-identification with the visual image of their body viewed from an out-of-body-perspective (Pfeiffer et al., 2014), ensuring that event encoding was considered self-relevant by the participants. Asynchronous (i.e., async) seen and felt brushstrokes reduce or abolish feelings of body ownership over the participant’s real body (in-body perspective) or invisible body below the cameras (out-of-body perspective). Thus, there were four experimental conditions: in-body sync, in-body async, out-of-body sync, out-of-body async. Seen and felt touches in the asynchronous condition were temporally spaced such that there was no overlap. Immediately after the induction of the embodiment illusion, the experimenter switched the video feed in the HMD unit to a live stream of the laboratory captured with the Ricoh Theta camera using a custom application implemented in Unity 5.3.0. The transition from recorded video to live stream was less than five seconds, ensuring a smooth transition.

#### Event Encoding

Immediately after each illusion induction, participants encoded interactive events selected to create realistic, distinct memories that could later be retrieved during functional scanning. The experimenter first opened the cabinets in front of the participant revealing stimuli to be used in the memory encoding stage of the experiment (see Figure 1C). Participants played four brief (i.e., less than 10 seconds) and emotionally neutral word games with the experimenter; “I spy”, “categories”, “sentence construction”, and the “truth game”. For the “I spy” game, the experimenter provided a colour (e.g., “I spy with my little eye something that is brown”) as a cue and participants made two guesses as to the identity of the object based on objects located in the right-hand cabinet (e.g., hat, cardboard box). In the “categories” game, the experimenter named a category (e.g., sports), prompting the participant to provide an example consistent with the category (e.g., cricket). The experimenter and participant took turns providing examples until a total of four were named. Each of the experimenter’s responses (e.g., weightlifting) was associated with a unique object (e.g., kettle bell) that was taken out of the left cabinet and held in front of the participant as it was named (see Table 1). During the “sentence construction” game, the participant and experimenter took two turns each verbalizing words to form a short sentence, beginning with the participant. The experimenter’s first response was associated with a unique object in the left cabinet to serve as a memory cue throughout the experiment, which was taken out of the cabinet and held in front of the participant as it was named (see Table 1). Lastly, the “truth game” involved the experimenter stating three consecutive statements based on a related topic, two of which were true and one of which was false (for full list of statements see Supplemental Table 1). Each statement was associated with a unique object in the left cabinet that was held in front of the participant as it was named (see Table 1). For example, the experimenter would select a toy bat and say “a group of bats is called a bevvy”, then select a toy owl and say “a group of owls is called a parliament”, then select a toy crow and say “a group of crows is called a murder” (see Figure 1C). The participant next guessed which statement was the lie (i.e., “a group of bats is called a bevvy”). The experimenter announced the game type (i.e., “I Spy”, “categories”,

**Table 1.**
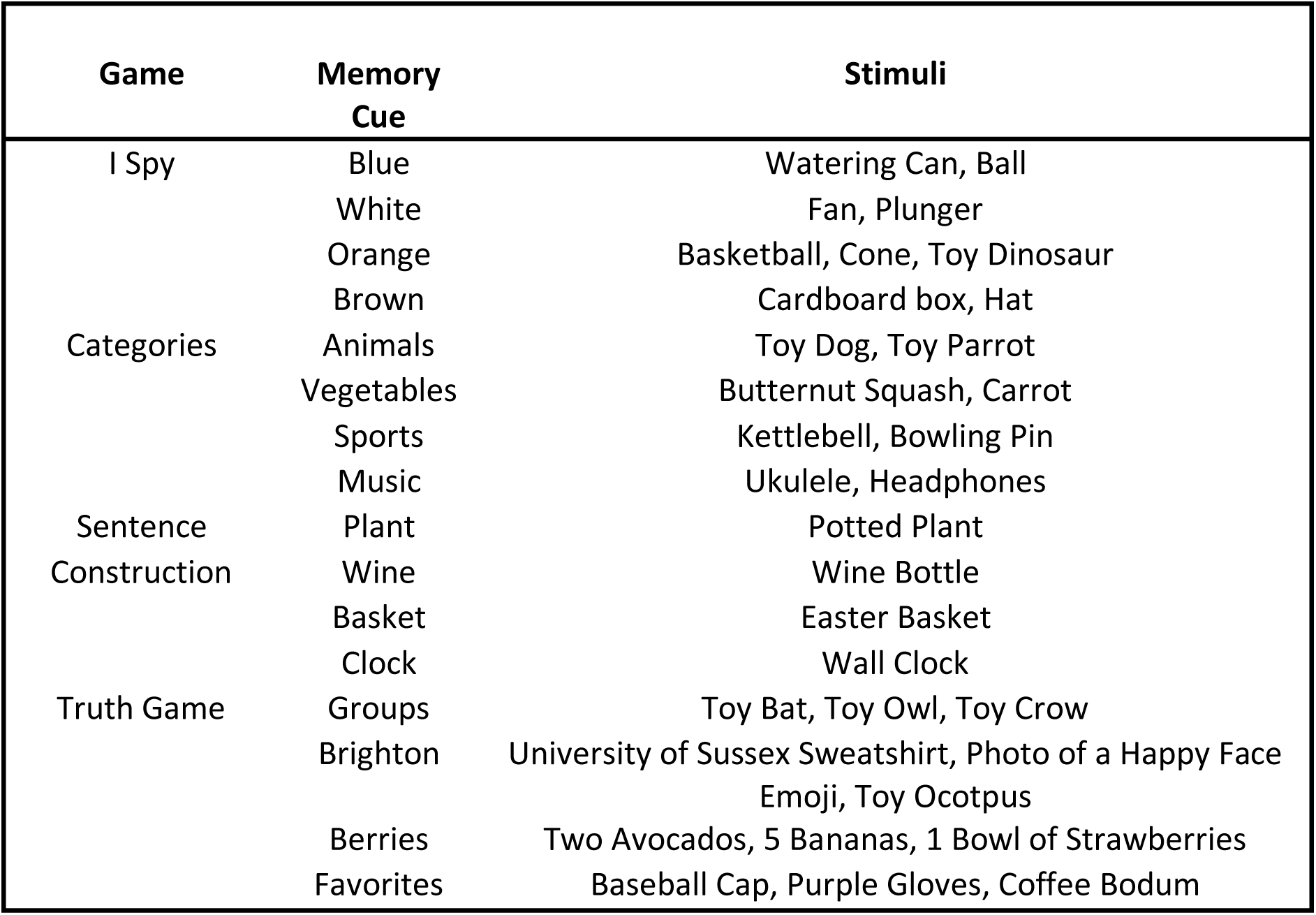
Word Game Stimuli.

“sentence construction2, and “truth game”) and a unique title (e.g. groups) at the start of each game. Each game was repeated using the exact same words a total of five consecutive times before beginning the next game. At the end of the last repetition, participants were instructed to close their eyes and retrieve the game from memory in as much detail as possible until the experimenter signalled them to stop after seven seconds, measured with a stopwatch. Repetitions and practiced retrieval were included to create a robust memory for each game that the participant would be able to retrieve from memory during functional scanning. While some games (e.g. I Spy) arguably required more detailed observation of the objects in the cabinet compared to others (e.g. sentence construction), differences between game types did not influence the behavioral or neuroimaging results since each of the four games was played in each of the four conditions. Thus, any differences between game types were the same across conditions.

Although no visuotactile stimulation was applied during the event encoding portion of the experiment, subjective feelings of body ownership induced by synchronous visuotactile stimulation persist up to five minutes after stimulation ends (Abdulkarim, Hayatou, & Ehrsson, 2021), which is roughly the amount of time it took to play the four word games in each condition. Thus, previous evidence indicates that the bodily illusions in the present experiment likely persisted throughout event encoding. Importantly, we assessed the effectiveness of the illusion induction at the end of event encoding period by asking participants to evaluate the bodily sensations they experienced when playing the word game tasks (see below).

#### Immediate Memory Test

After each round of four games, participants were instructed to remove the HMD unit and answered a series of questions. The first was a questionnaire designed to assess bodily sensations experienced during the illusion induction and while playing the games, adapted from Guterstam and colleagues (2015a). It included three statements designed to assess the strength of the embodiment illusion (see Table 2, I1 to I3) and three control statements to assess a participant’s susceptibility to demand characteristics (see Table 2, C1 to C3) that were rated on a 7-point Likert scale from –3 (i.e., strongly disagree) to 0 (i.e., neutral) to 3 (i.e., strongly agree), with 0 reflecting neutral.

**Table 2.**
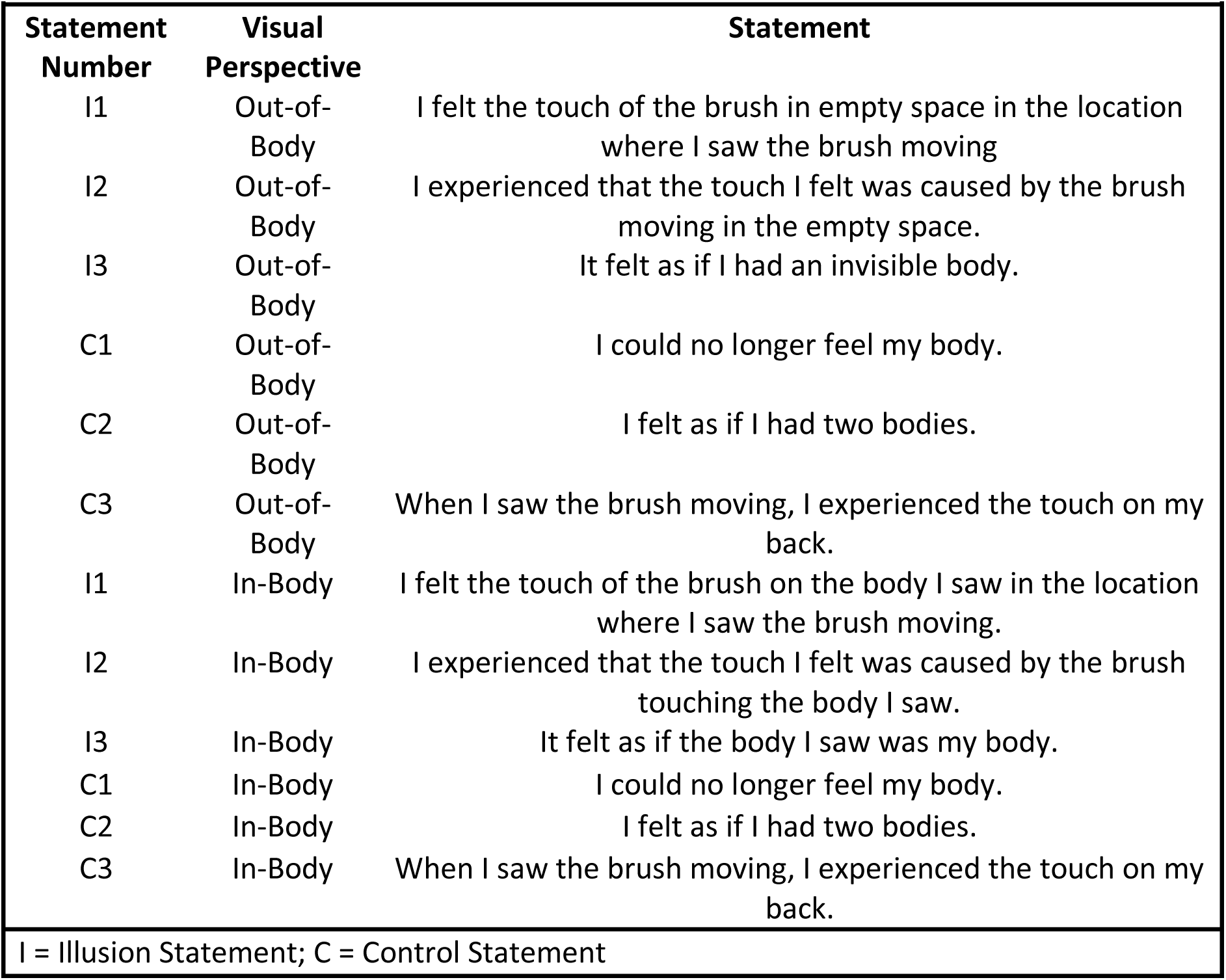
Bodily Illusion Questionnaire Items.

| Statement Number | Visual Perspective | Statement |
| --- | --- | --- |
| I1 | Out-of-Body | I felt the touch of the brush in empty space in the location where I saw the brush moving |
| I2 | Out-of-Body | I experienced that the touch I felt was caused by the brush moving in the empty space. |
| I3 | Out-of-Body | It felt as if I had an invisible body. |
| C1 | Out-of-Body | I could no longer feel my body. |
| C2 | Out-of-Body | I felt as if I had two bodies. |
| C3 | Out-of-Body | When I saw the brush moving, I experienced the touch on my back. |
| I1 | In-Body | I felt the touch of the brush on the body I saw in the location where I saw the brush moving. |
| I2 | In-Body | I experienced that the touch I felt was caused by the brush touching the body I saw. |
| I3 | In-Body | It felt as if the body I saw was my body. |
| C1 | In-Body | I could no longer feel my body. |
| C2 | In-Body | I felt as if I had two bodies. |
| C3 | In-Body | When I saw the brush moving, I experienced the touch on my back. |

I = Illusion Statement; C = Control Statement

Next, participants answered one cued recall question from each game (see Supplemental Table 2), and rated their memory for each game on 7-point subjective ratings according to vividness (1 = a little, 7 = a lot), emotional intensity (1 = a little, 7 = a lot), perceived accuracy (i.e., the degree to which participants felt their memory was an accurate representation of the game; 1 = a little, 7 = completely accurate), and visual perspective separately for first-person perspective (1PP) and third-person perspective 3PP ratings (1 = a little, 7 = exclusively 1PP/3PP). Participants were instructed that 1PPs referred to visualizing their mental scene from the perspective of their own eyes such that they did not see themselves in the event, whereas 3PPs referred to being able to see themselves in the mental scene as if from the perspective of an observer. Thus, 1PP ratings correspond to the tendency to retrieve events from an in-body visual perspective and 3PP ratings correspond to the tendency to retrieve events from an out-of-body visual perspective. We use separate terminology for the ratings to distinguish them from our experimental manipulation of visual perspective. The 1 to 7 scale during the immediate memory test was used to align with previous research using 7-point Likert scales (e.g., Iriye & St. Jacques, 2021).

Once participants had answered all questions, the next embodiment illusion was induced. This process was repeated for each of four conditions: in-body sync, in-body async, out-of-body sync, out-of-body async. Thus, participants played four games in each condition. The order of conditions, games, and stimuli within each game were allocated randomly for each participant. The behavioral session, including video recording and editing, the illusion inductions, event encoding, and the cued recall test and subjective ratings, took approximately two hours per participant to complete.

#### fMRI Scanning

Later that day (i.e., 1.5 to 3 hours following memory encoding), participants retrieved memories for each game during functional scanning. Participants were free to spend the time between testing sessions as they wished. Before scanning, participants were shown the title of each memory and asked to report the associated game and stimuli to ensure they were able to recall each event using the memory cue. All participants were able to recall each event. Next, participants underwent a practice session to familiarize themselves with the task and timing of responses, which involved retrieving their memory for each event once and making subjective ratings of visual perspective and vividness.

Scanning commenced after completion of the practice session. On each trial, participants were presented with the memory cue (e.g., groups) and game title (e.g., truth game). This prompt was quickly followed (i.e., 800 ms) by an instruction to close their eyes, at which point they were asked to retrieve the memory for the event in as much detail as possible until an auditory tone sounded through MRI-compatible headphones 6.4 s later. Upon hearing the brief auditory tone, participants were instructed to stop retrieving the event and open their eyes. They were then asked to provide subjective ratings of the degree to which they retrieved the event from an 1PP, the degree to which they retrieved the event from an 3PP, and the degree of vividness associated with memory retrieval, each on four-point scales from one = low to four = high. We adopted a 1 to 4 scale for the in-scanner ratings as the only MR-compatible button box available had four buttons. Note that we used a 7-point Likert scale for the 1PP/3PP ratings in the immediate memory test to align with previous research (e.g., Iriye & St. Jacques, 2021). However, our goal was to establish whether encoding events from an in-body perspective led to higher 1PP ratings and encoding events form an out-of-body perspective led to higher 3PP ratings, not to compare changes in 1PP/3PP ratings between sessions. Accordingly, the different scales do not impact the interpretation of our results. The order of the 1PP and 3PP ratings was counterbalanced across participants to control for potential order effects. Participants had 2.4 s for each rating and responded using a four button MRI-compatible response box. An example trial is depicted in Figure 1D.

There were 12 functional runs consisting of 16 trials (i.e., one trial per event, four trials per condition), resulting in a total of 48 trials per condition. Trial order was randomized for each functional run. Trials were separated by a jittered fixation cross, which was equally spaced across a variable length (i.e., 1.6 to 8 s) and distributed exponentially such that shorter inter-trial intervals occurred more frequently than longer intervals.

#### Post-Scanning Memory Test

Immediately after scanning, participants answered a cued recall question different from the one asked immediately following memory encoding (see Appendix B) and made subjective ratings of emotional intensity and perceived memory accuracy. fMRI scanning and the post-scanning memory test together lasted 2.5-3 hours per participant.

### MRI Data Acquisition and Preprocessing

Functional and structural images were collected on a 3T MAGNETOM Prisma MRI scanner. Detailed anatomical data were collected using a multi-planar rapidly acquired gradient echo (MPRAGE) sequence. Functional images were acquired using a T2*-weighted echo planar sequence (TR = 800 ms, TE = 37 ms, FOV = 208 x 208 mm, Slice Thickness = 2 mm). Whole brain coverage was obtained via 72 interleaved slices, acquired at an angle corresponding to AC-PC alignment, with a multiband factor of 8 and a 2 mm x 2 mm in-plane resolution. The first ten volumes of each run were discarded to allow for T1 equilibrium.

Preprocessing of functional images was performed using SPM12. Functional images were realigned within and across runs to correct for head movement, segmented into gray matter, white matter, and cerebrospinal fluid, co-registered to the participant’s anatomical image, and spatially normalized to the Montreal Neurological Institute (MNI) template. BOLD signal response patterns for each trial were estimated using a general linear model (GLM). A regressor was estimated for each trial in the twelve functional runs, resulting in 16 beta estimates per run (i.e., 4 trials per condition per run; 48 trials per condition in total).

Regressors were time-locked to the onset of the memory cue and the duration set to cover the memory retrieval period (i.e., 6.4s), excluding the auditory tone and subjective ratings. Six movement parameters were included as separate regressors. The SPM canonical haemodynamic response basis function was used to estimate brain responses. Additionally, GLM’s were estimated using the fast serial correlations option to account for the multiband sequence employed during neuroimaging data collection. The functional data was denoised using the RobustWLS Toolbox (Diedrichsen & Shadmehr, 2005).

### Behavioral Analyses

The illusion induction questionnaire data was assessed by submitting average illusion minus control statement scores for each condition (e.g., Iriye et al., 2022; 2024) to a 2 (visual perspective: in-body, out-of-body) x 2 (embodiment: sync, async) repeated-measures ANOVA. Illusion minus control statement scores reflect the strength of the illusion induction, taking into account participants’ susceptibility to answer positively to the control statements. 1PP ratings, 3PP ratings, vividness ratings, emotional intensity ratings, and belief in memory accuracy ratings from the immediate memory test, scanning, and post-scanning memory test were assessed with eight separate 2 (visual perspective: in-body, out-of-body) x 2 (embodiment: sync, async) repeated-measures ANOVAs. We controlled for multiple comparisons using Bonferroni-Holm corrections. Frequentist statistics were supported with Bayesian analyses. Outliers were defined as data points that were 1.5 interquartile ranges below the first quartile or above the third quartile. All analyses were conducted with and without outliers to determine their influence on the results. All major conclusions remain the same. Data with outliers is reported in the main text, while data without outliers is reported in the Supplemental Material. See Supplemental Table 3 for the means and standard deviations of the immediate memory test, scanning, and post-scanning memory tests not reported in the main text. All behavioral analyses were analyzed in JASP version 0.19 (JASP Team, 2024).

### Multivariate Decoding

#### ROI Analyses: Classifier Accuracy Across Conditions

We created individual masks of the left and right hemispheres of angular gyrus, precuneus, and hippocampus in WFU Pickatlas (Maldjian et al., 2003) based on the Individual Brain Atlases using Statistical Parametric Mapping (IBASPM; Alemán-Gómez, Melie-García, Valdés-Hernandez, 2006).

Masks were then converted into binary format using the MarsBar toolbox for SPM (Brett et al., 2002) and resampled to 2mm cubic voxels to match the dimension of the beta estimates obtained from the GLM analysis (see Figure 2).

**Figure 2.**
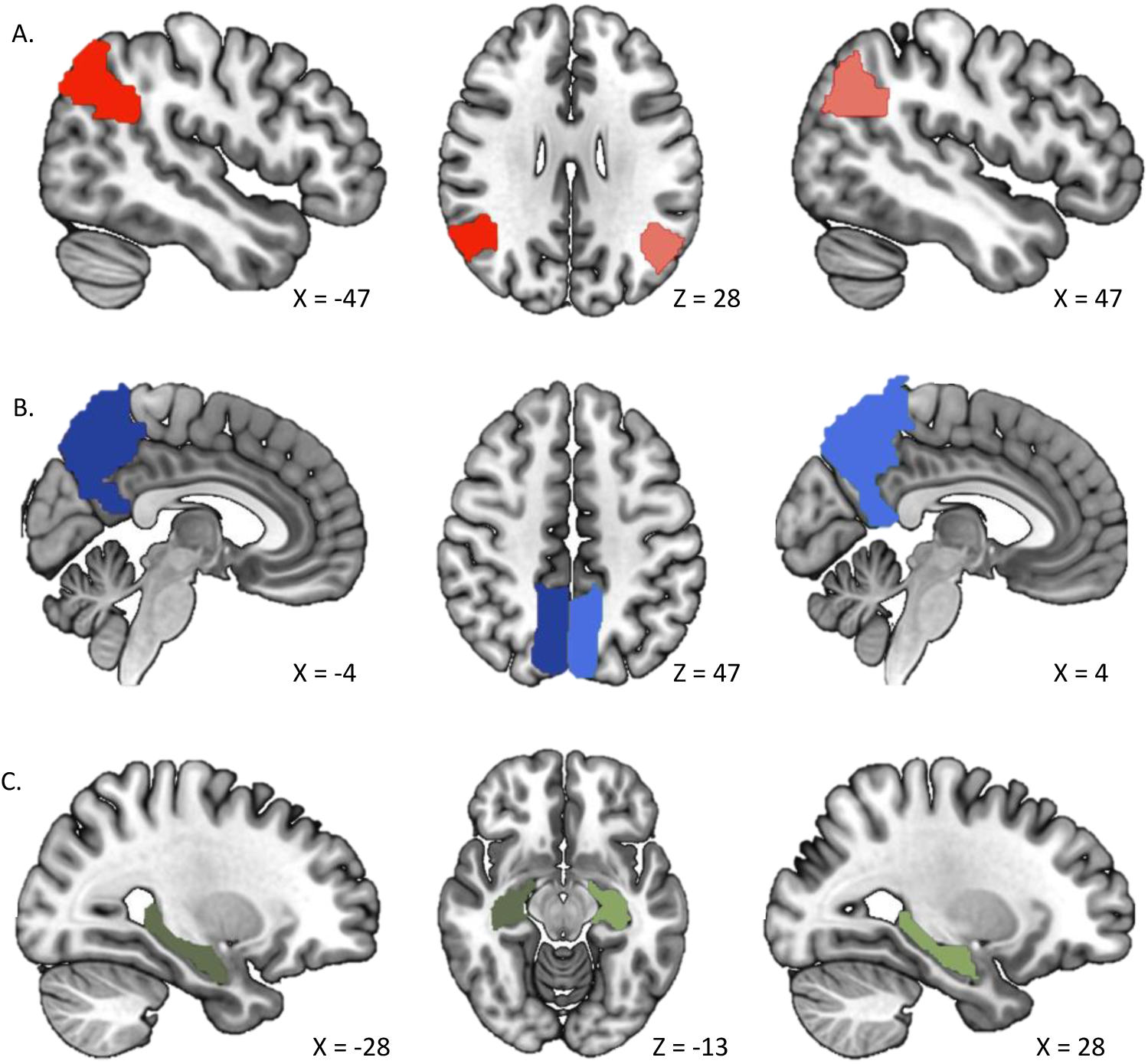
ROIs for the left and right angular gyrus. (A), precuneus (B), and hippocampus (C) as ROIs. Separate masks were created for the left and right hemispheres of each region.

Two separate decoding analyses assessed whether patterns of neural activity in each of the six ROIs predicted 1) the visual perspective initially experienced during encoding (i.e., in-body vs. out-of-body) and 2) feelings of embodiment initially experienced during encoding (i.e., sync versus async visuotactile stimulation). Next, we conducted an additional two analyses to determine whether any of the six ROIs responded to the interaction between visual perspective and embodiment by holding embodiment constant while varying visual perspective. One compared patterns of activity in the in-body sync compared to out-of-body sync conditions (i.e., decoding visual perspective for events encoded with strong embodiment). The other contrasted the in-body async with the out-of-body async condition (i.e., decoding visual perspective for events encoded with weak embodiment). We compared these two analyses by investigating whether each of the six ROIs could decode visual perspective for memories encoded with both sync and async visuotactile stimulation, either sync or async visuotactile stimulation, or neither. The comparison of these two analyses revealed whether patterns of activity within a given ROI differentiated between visual perspectives for memories associated with strong and weak embodiment separately, which would indicate the integration of visual perspective and embodiment within memories of past events.

For each of the four ROI analyses, we calculated the decoding accuracy (i.e., number of correctly classified trials / number of total trials) for each participant within each ROI using a linear discriminant analysis (LDA) classifier. The data was divided into training and test sets using k-fold cross-validation. Subject-level accuracies were submitted to a two-sided, one-sample t-test against a null hypothesis of chance-level classification accuracy (i.e., 50%). We corrected for multiple comparisons by applying Bonferroni-Holm corrections. All multivariate analyses were implemented in the CoSMoMVPA toolbox (Oosterhof et al., 2016). We calculated Bayes factors to supplement frequentist statistics in JASP verson 0.19 (JASP Team, 2024). Crucially, the neuroimaging results relate to the altered feelings of bodily selfhood induced by the illusions, and not low-level discrepancies in visuotactile synchronicity between the sync and async conditions, since no visuotactile stimulation was present during encoding of the word games which participants repeatedly retrieved during fMRI scanning.

We performed a supplementary decoding analysis testing whether patterns of activity in each of the ROIs could predict the specific game associated with a given trial as certain game types may engage visual perspective and embodiment more than others (e.g., I Spy more so than Sentence Construction), which may have affected overall classifier accuracy in the main decoding analyses. For each ROI, we calculated a classification accuracy score per game and compared them with a one-way repeated-measures ANOVA to investigate whether classifier performance varied between game types (see Supplemental Material).

A limitation of the ROI approach is that it assumes that the pre-specified regions are functionally homogenous, which may overlook the complex and varied functional architecture of the brain (Poldrack, 2007). A separate concern is that focusing on specific regions may miss important activity patterns in other parts of the brain (Poldrack, 2007).

#### Whole-brain searchlight analyses

We repeated each of the four main decoding analyses described above (i.e., in-body vs. out-of-body, sync vs. async, sync in-body vs. sync out-of-body, async in-body vs. async out-of-body) at the whole-brain level to investigate whether any additional brain regions outside of the pre-specified ROIs could predict visual perspective, embodiment, or their interaction. For each subject, a sphere comprised of 100 voxels was fitted around each voxel in the acquired volumes to create searchlight maps for each participant that reflected the classification accuracies determined by k-fold cross-validation estimated by a LDA classifier. Chance-level accuracy (i.e., 50%) was subtracted from each voxel in each participant’s classification accuracy map. We investigated group-level effects by submitting the resulting subject-specific classification accuracy maps to a one-sample t-test against a null hypothesis of chance-level accuracy (i.e., 0). The resulting group-level classification accuracy map was thresholded at *p_FWE_* < .05 to correct for multiple comparisons.

While the whole brain searchlight approach addresses limitations of the ROI analyses, it comes with its own caveats. Searchlight analyses involve numerous statistical tests across the entire brain requiring stringent corrections for multiple comparisons, which can distort the data with both false positive and false negative results (Etzel, Zacks, & Braver, 2013). To address this issue, we report the results thresholded at the less conservative threshold of *p* < .01 and cluster size > 5 voxels in the Supplemental Material.

We used MRIcroGL (v1.2.20220720) to create 3D surface visualizations of the results and project the statistical maps onto individual slices onto the MNI152 T1-weighted template. Identified regions in each of the analyses were anatomically labelled by cross-referencing the co-ordinates with Duvernoy’s brain atlas (Duvernoy, 1999). By implementing both ROI and whole-brain searchlight analyses, we aimed to provide a balanced and comprehensive account of regions involved in memory retrieval of events formed from different visual perspectives and feelings of embodiment.

## Results

### Behavioral Results

#### Event encoding: Illusion Induction Questionnaire

A 2 (Embodiment: sync, async) x 2 (Visual Perspective: in-body, out-of-body) repeated measures ANOVA on average illusion minus control statement ratings found main effects of both embodiment, *F* (1, 23) = 24.35, *p* < .001, η_p_2 = .51, BF_10_ = 390.38, and visual perspective, *F* (1, 23) = 21.66, *p* < .001, η_p_2 = .49, BF_10_ = 103.03. There was also an interaction between embodiment and visual perspective, *F* (1, 23) = 5.40, *p* = .029, η_p_2 = .19, BF_10_ = 138 771.60. Follow-up pairwise t-tests confirmed that illusion minus control statement ratings were higher in the synchronous compared to asynchronous conditions for both in-body, *t* (23) = 4.69, *p* = .004, Cohen’s d = 1.34, and out-of-body perspectives, *t* (23) = 3.35, *p* = .008, Cohen’s d = .69 (see Figure 3). Bayes factors confirmed the difference between synchronous and asynchronous illusion minus control statement ratings for the in-body perspective (BF_10_ = 3844.05), but suggested this difference was not as strong for the out-of-body perspective (BF_10_ = .90). For means and standard deviations for each condition, see Table 3. See Supplemental Figure 1 for the results of individual questionnaire items.

**Figure 3.**
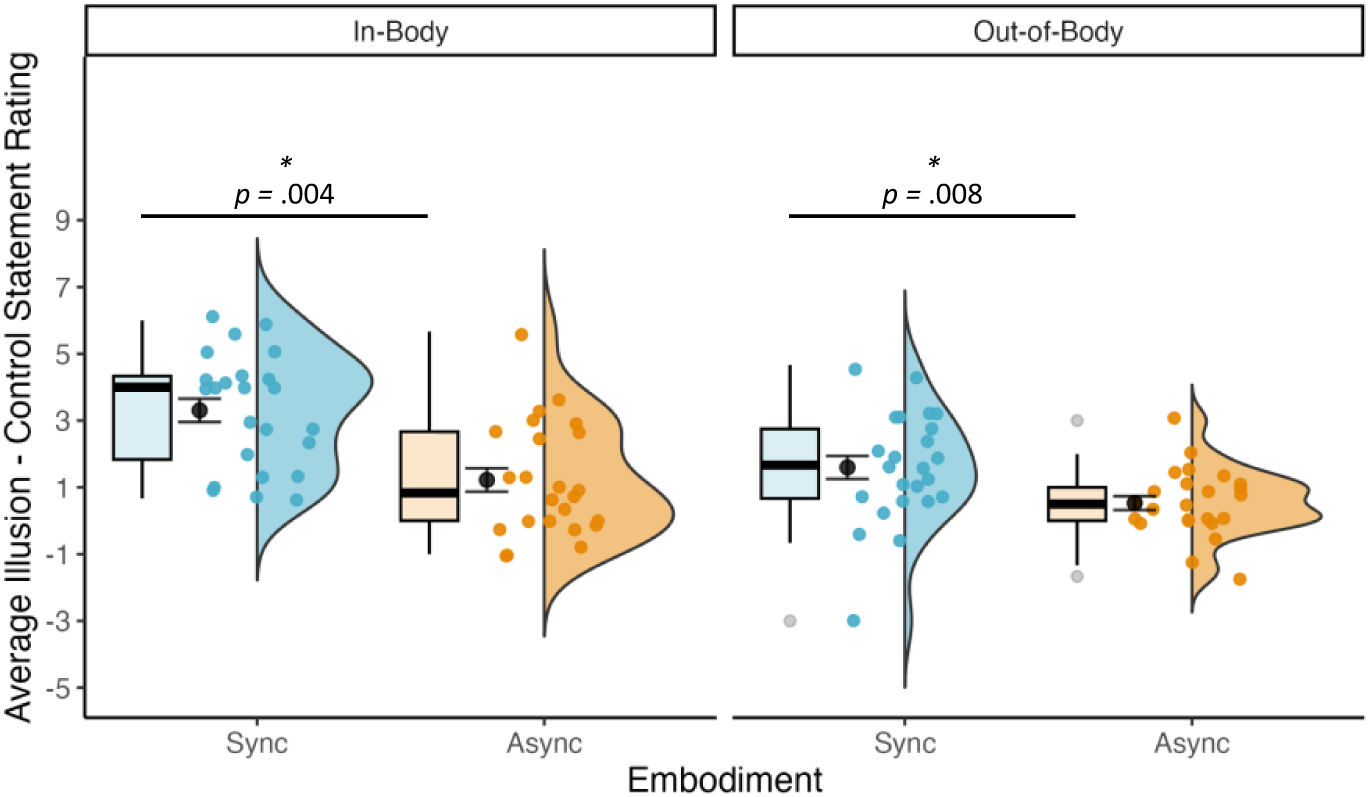
Average illusion minus control statement ratings were higher in the synchronous compared to asynchronous conditions for both in-body and out-of-body perspectives, indicating that the illusion worked as expected. Sync = synchronous, Async = asynchronous. 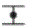 = mean +/− standard error.

**Table 3.**
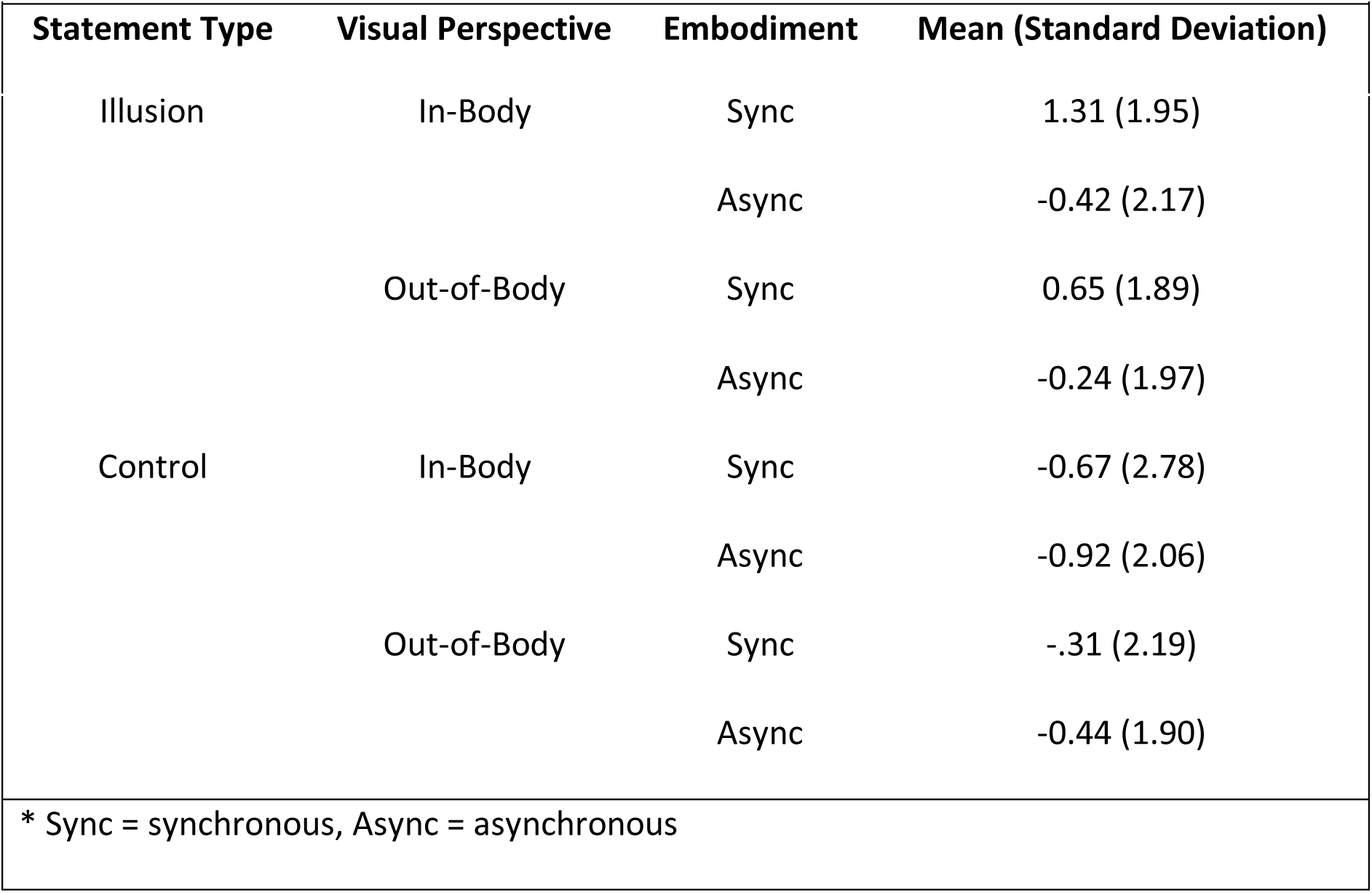
Illusion Induction Questionnaire Results.

| Statement Type | Visual Perspective | Embodiment | Mean (Standard Deviation) |
| --- | --- | --- | --- |
| Illusion | In-Body | Sync | 1.31 (1.95) |
|  |  | Async | -0.42 (2.17) |
|  | Out-of-Body | Sync | 0.65 (1.89) |
|  |  | Async | -0.24 (1.97) |
| Control | In-Body | Sync | -0.67 (2.78) |
|  |  | Async | -0.92 (2.06) |
|  | Out-of-Body | Sync | -.31 (2.19) |
|  |  | Async | -0.44 (1.90) |
| * Sync = synchronous, Async = asynchronous |  |  |  |

#### Immediate memory test

A 2 x 2 repeated measures ANOVA with visual perspective (in-body, out-of-body) and embodiment (sync, async) conducted on average cued recall accuracy scores did not reveal a main effect of visual perspective, *F*(1,23) = .49, *p* = .49, η_p_2 = .02, BF_10_ = 0.33, embodiment, *F*(1,23) < .001, *p* = 1.00, η_p_2 < .001, BF_10_ = 0.27, or an interaction, *F*(1,23) = 2.09, *p* = .16, η_p_2 = .08, BF_10_ = .09. Thus, immediate memory recall was equal between conditions. To investigate whether the visual perspective participants adopted at immediate retrieval differed according to condition, we conducted a 2 (visual perspective: in-body, out-of-body) x 2 (embodiment: sync, async) repeated measures ANOVA for 1PP and 3PP ratings separately. The analysis of 1PP ratings revealed an expected main effect of visual perspective, *F* (1,23) = 57.98, *p* < .001, η_p_2 = .72, BF_10_ = 168 580.80, where 1PP ratings were higher in the in-body (*M* = 5.86, *SD* = 1.35) compared to out-of-body (*M* = 3.77, *SD* = 1.66) perspective (see Figure 4A). There was no significant main effect of embodiment, *F* (1,23) = 2.53, *p* = .13, η_p_2 = .10, BF_10_ = .70, or visual perspective x embodiment interaction, *F* (1,23) = 0.16, *p* = .70, η_p_2 = .01, although a Bayesian analysis indicated very strong evidence that a model including the model was better at explaining the data than the null model (BF_10_ = 34502.70). For the 3PP ratings, there was a significant main effect of visual perspective, *F* (1,23) = 70.49, *p* < .001, η_p_2 = .75, BF_10_ = 940 560.59 indicating higher 3PP ratings in out-of-body (*M* = 4.01, *SD* = 1.85) compared to in-body (*M* = 1.74, *SD* = 1.20) perspectives (see Figure 4B). There was no main effect of embodiment, *F* (1,23) = .52, *p* = .48, η_p_2 = .02, BF_10_ = .31. The interaction between visual perspective and embodiment was significant, *F*(1,23) = 5.53, *p* = .03, η_p_2 = .19, BF_10_ = 811 963.71, but did not reveal differences between synchronous and asynchronous visuotactile stimulation in either the in-body (sync: *M* = 1.62, *SD* = 1.10; async: *M* = 1.87, *SD* = 1.09, BF_10_ = 1.22) or out-of-body (sync: *M* = 4.27, *SD* = 1.52; async: *M* = 3.75, *SD* = 1.80, BF_10_ = .60) perspective, *p*’s > .09.

**Figure 4.**
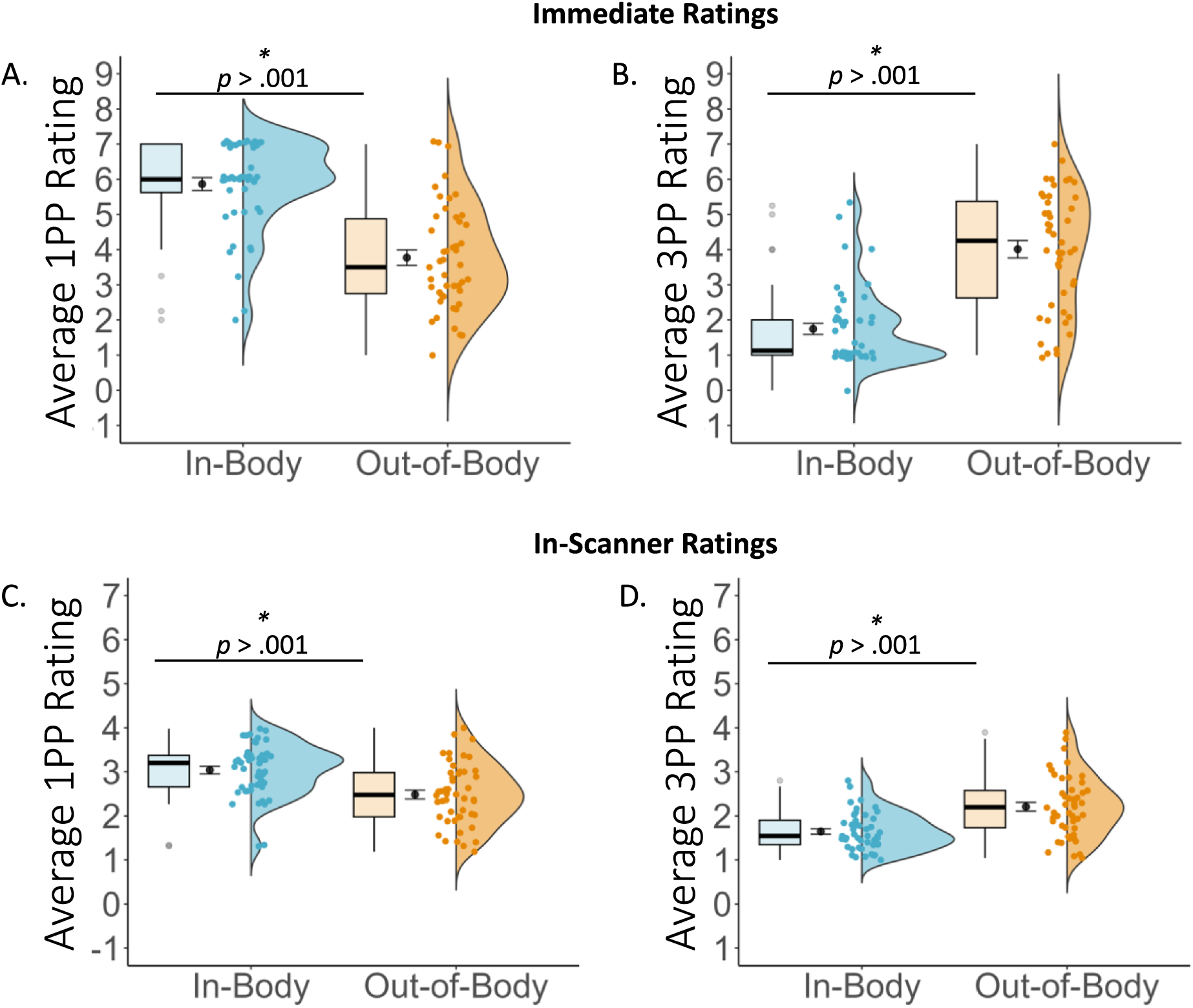
As expected, average 1PP ratings were higher in the in-body perspective. (A), whereas average 3PP ratings were higher in the out-of-body perspective (B). Similarly, average in-scanner 1PP ratings were higher in the in-body perspective (C), whereas average in-scanner 3PP ratings were higher in the out-of-body perspective (D). 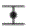 = mean +/− standard error.

For the vividness ratings, a 2 (visual perspective: in-body, out-of-body) x 2 (embodiment: sync, async) did not find a main effect of visual perspective, *F*(1,23) = .25, *p* = .62, η_p_2 = .01, BF_10_ = .38, embodiment, *F*(1,23) = .34, *p* = .56, BF_10_ = .28, η_p_2 = .02, or interaction, *F*(1,23) = .05, *p* = .83, η_p_2 = .002, BF_10_ = .07. Similarly, there was was no main effect of visual perspective, *F* (1,23) = 2.39, *p* = .14, η_p_2 = .10, BF_10_ = .61, embodiment, *F* (1,23) = .04, *p* = .84, η_p_2 = .002, BF_10_ = .28, or interaction, *F* (1,23) = 1.96, *p* = .18, η_p_2 = .08, BF_10_ = .14, for the emotional intensity ratings. The same was the case for the belief in memory accuracy ratings (visual perspective: *F* (1,23) = 2.28, *p* = .14, η_p_2 = .09, BF_10_ = .67; embodiment: *F* (1,23) = 2.06, *p* = .16, η_p_2 = .08, BF_10_ = .58; interaction: *F* (1,23) = .33, *p* = .57, η_p_2 = .01, BF_10_ = .13).

Together, the results indicate that the visual perspective manipulation was successful such that participants recalled whether events were experienced from an in-body or out-of-body perspective directly after encoding memories for the word games. Importantly, there were no significant differences between the conditions for memory phenomenology, such that vividness, emotional intensity, and perceived memory accuracy were equal across the conditions.

#### In-scanner subjective ratings

The analysis of in-scanner visual perspective ratings revealed the same pattern of effects observed in the immediate memory test (see Figure 4C&D). For the 1PP ratings, there was a significant effect of visual perspective, *F* (1,23) = 26.25, *p* < .001, η_p_2 = .53, BF_10_ = 167 684.53, indicating higher 1PP ratings in the in-body (*M* = 3.05, *SD =* .97) compared to out-of-body perspective conditions (*M* = 2.49, *SD* = 1.07). There was no main effect of embodiment, *F* (1,23) = 1.02, *p* = .32, η_p_2 = .04, BF_10_ = .67, or interaction effect, *F* (1,23) < .001, *p* = .97, η_p_2 < .001, although a Bayesian analysis indicated very strong evidence that a model including the interaction term was better at predicting the data compared to the null model (BF_10_ = 33 227.60). For the 3PP ratings, there was also a significant effect of visual perspective, *F* (1,23) = 25.21, *p* < .001, η_p_2 = .52, BF_10_ = 899 769.65. 3PP ratings were higher for the out-of-body perspective conditions (*M* = 2.22, *SD* = 1.11) compared to the in-body conditions (*M* = 1.65, *SD* = .88). There was no main effect of embodiment, *F* (1,23) = .68, *p* = .42, η_p_2 = .03, BF_10_ = .33, or interaction effect, *F*(1,23) = .006, *p* = .94, η_p_2 < 0.001, although a Bayesian analysis indicated very strong evidence that a model including the interaction term was better at predicting the data compared to the null model (BF_10_ = 896 866.40). Turning to the vividness ratings, there was no main effect of visual perspective, *F* (1,23) = .65, *p* = .43, η_p_2 = .03, BF_10_ = .58, or embodiment, F(1,23) = 2.48, *p* = .13, η_p_2 = .10, BF_10_ = .39, or an interaction, *F*(1,23) = 61, *p* = .44, η_p_2 = .03, BF_10_ = .09. These results indicate that participants could remember the visual perspective events were initially formed from during retrieval, while the vividness of these memories remained consistent across the different conditions.

#### Post-scanning cued recall and subjective ratings

The analysis of cued recall accuracy revealed a marginally significant effect of visual perspective, *F* (1,23) = 4.39, *p* = .05, η_p_2 = .16, BF_10_ = .44, which was qualified by an interaction between visual perspective and embodiment n, *F* (1,23) = 5.37, *p* = .03, η_p_2 = .19, BF_10_ = 2.09. A follow-up post-hoc test revealed that memory accuracy was higher in the in-body sync condition compared to the out-of-body sync condition, *p* = .02, BF_10_ = 5.22 (Table 3). None of the other post-hoc comparisons were significant, all *p*’s > .13.

Turning to the emotional intensity ratings, there was no main effect of visual perspective, *F* (1,23) = .74, *p* = .40, η_p_2 = .03, BF_10_ = .32, embodiment, *F* (1,23) = .05, *p* = .83, η_p_2 = .002, BF_10_ = .26, or interaction, *F* (1,23) = .05, *p* = .83, η_p_2 = .002, BF_10_ = .07. Similarly for the belief in memory accuracy ratings, there was no main effect of visual perspective, *F* (1,23) = 1.97, *p* =.17, η_p_2 = .08, BF_10_ = .60, embodiment, *F*(1,23) = .57, *p* = .46, η_p_2 = .22, BF_10_ = .34, or interaction, *F* (1,23) = 1.61, *p* = .22, η_p_2 = .07, BF_10_ = .12.

**Table 3.**
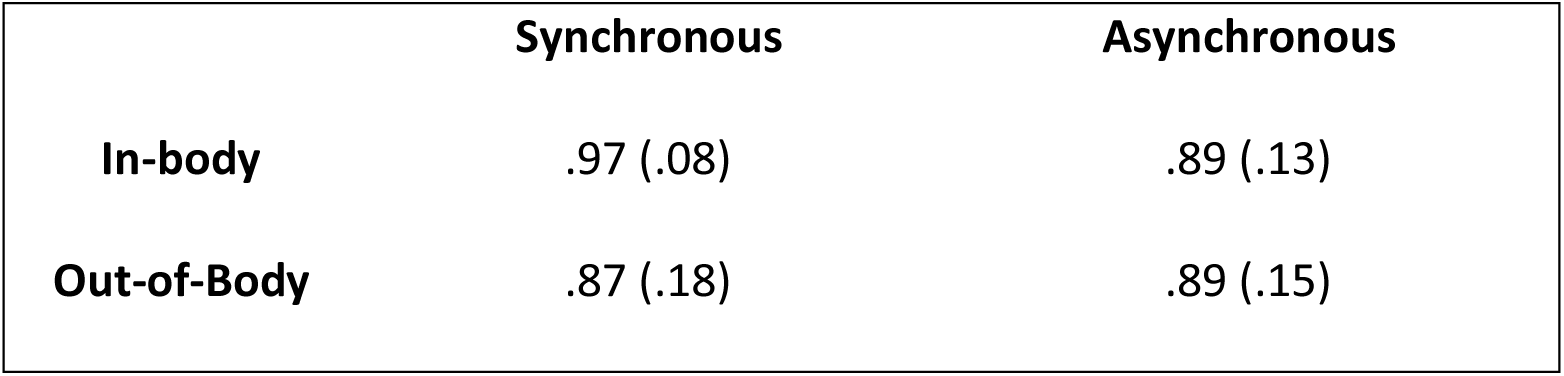
Post-Scanning: Mean cued recall accuracy scores (standard deviation)

|  | Synchronous | Asynchronous |
| --- | --- | --- |
| **In-body** | .97 (.08) | .89 (.13) |
| **Out-of-Body** | .87 (.18) | .89 (.15) |

Thus, average illusion statement scores were greater following sync compared to async visuotactile stimulation for events encoded from an in-body compared to out-of-body perspectives. Post-scanning cued recall accuracy was higher in the in-body perspective compared to out-of-body perspective only in the synchronous conditions. Participants successfully recalled the visual perspective immediately after memory encoding and during functional scanning. However, memory vividness, emotional intensity, and perceived memory accuracy were the same across conditions at both testing sessions.

### Multivariate Decoding Analyses Results: ROI Analyses

#### Visual perspective

Frequentist statistics indicated that patterns of activity during retrieval did not predict the visual perspective (i.e., in-body, out-of-body) of a given trial significantly greater than chance in any of the ROIs after applying Bonferroni-Holm corrections to account for multiple comparisons (see Figure 5). However, a Bayesian analysis revealed anecdotal evidence in favor of greater than chance decoding accuracies in the precuneus (left: BF_10_ = 2.76; right: BF_10_ = 1.09) and left angular gyrus (BF_10_ = 1.26). We observed moderate evidence against greater than chance decoding accuracies in the right angular gyrus (BF_10_ = 0.26), and hippocampus (left: BF_10_ = 0.28; right: BF_10_ = 0.33).

**Figure 5.**
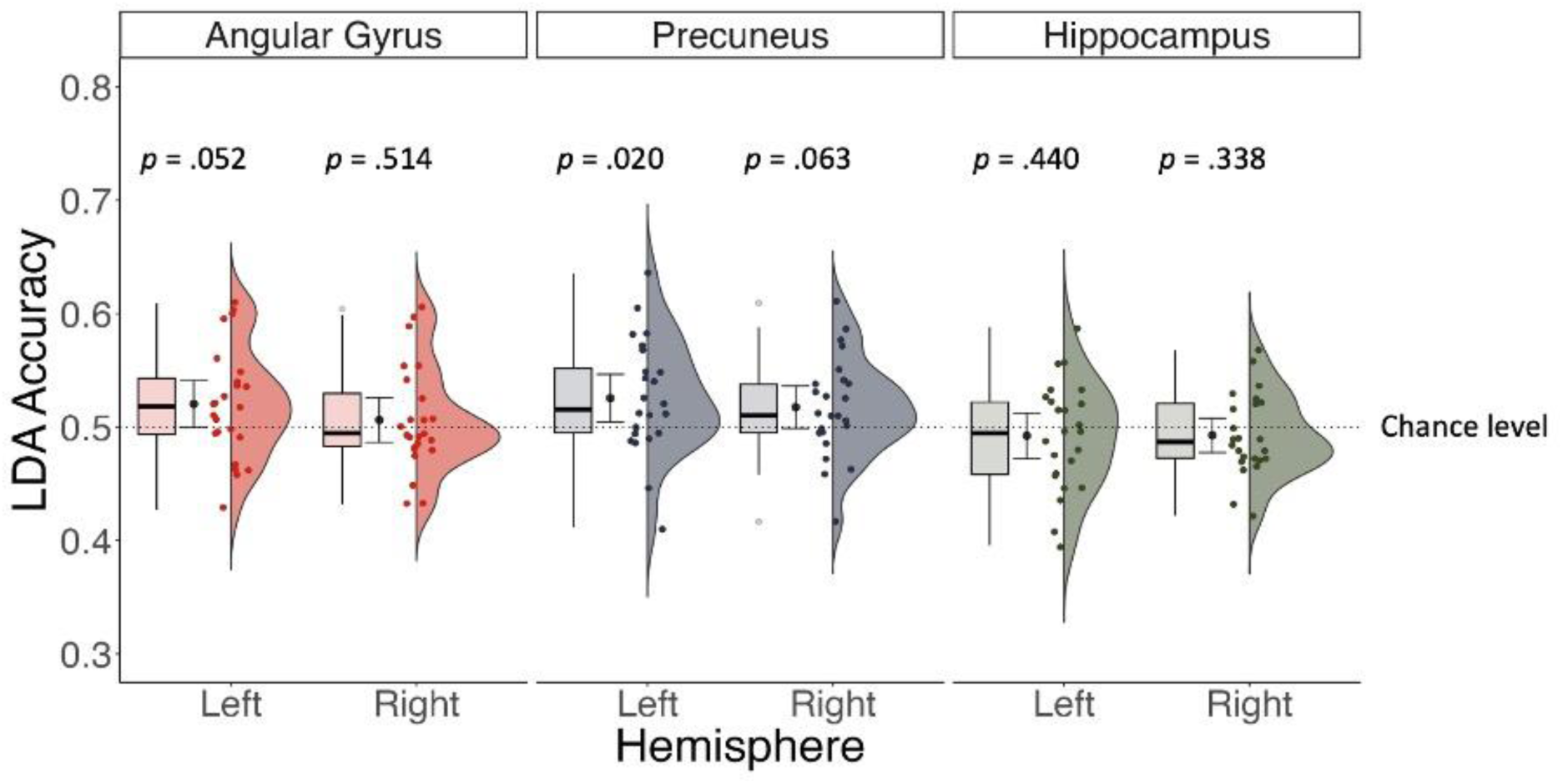
Frequentist statistics indicated decoding accuracy was no different from chance in any of the ROIs when predicting visual perspective based on patterns of activity during memory retrieval after correcting for multiple comparisons. However, Bayesian results indicated weak evidence supporting above chance decoding accuracy in the left angular gyrus and bilateral precuneus. 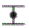 = mean decoding accuracy +/− 95% confidence interval.

#### Embodiment

Patterns of voxel activation during retrieval in both the left angular gyrus, *t*(23) = 3.965, *p* < .001, Cohen’s d = 0.791, BF_10_ = 53.901 (*M* = .529, *SD* = .035), left precuneus, *t*(23) = 3.83, *p* < .001, Cohen’s d = .787, BF_10_ = 40.229 (*M* = .537, *SD* = .047), and right precuneus, *t*(23) = 3.277, *p* = .003, BF_10_ = 12.404, Cohen’s d = .674 (*M* = .531, *SD* = .046), distinguished the embodiment associated with memory encoding (see Figure 6). A Bayesian analysis indicated weak evidence against greater than chance decoding accuracies in the left hippocampus (BF_10_ = .393) and moderate evidence against greater than chance decoding accuracies in the right hippocampus (BF_10_ = .233) and right angular gyrus (BF_10_ = .233).

**Figure 6.**
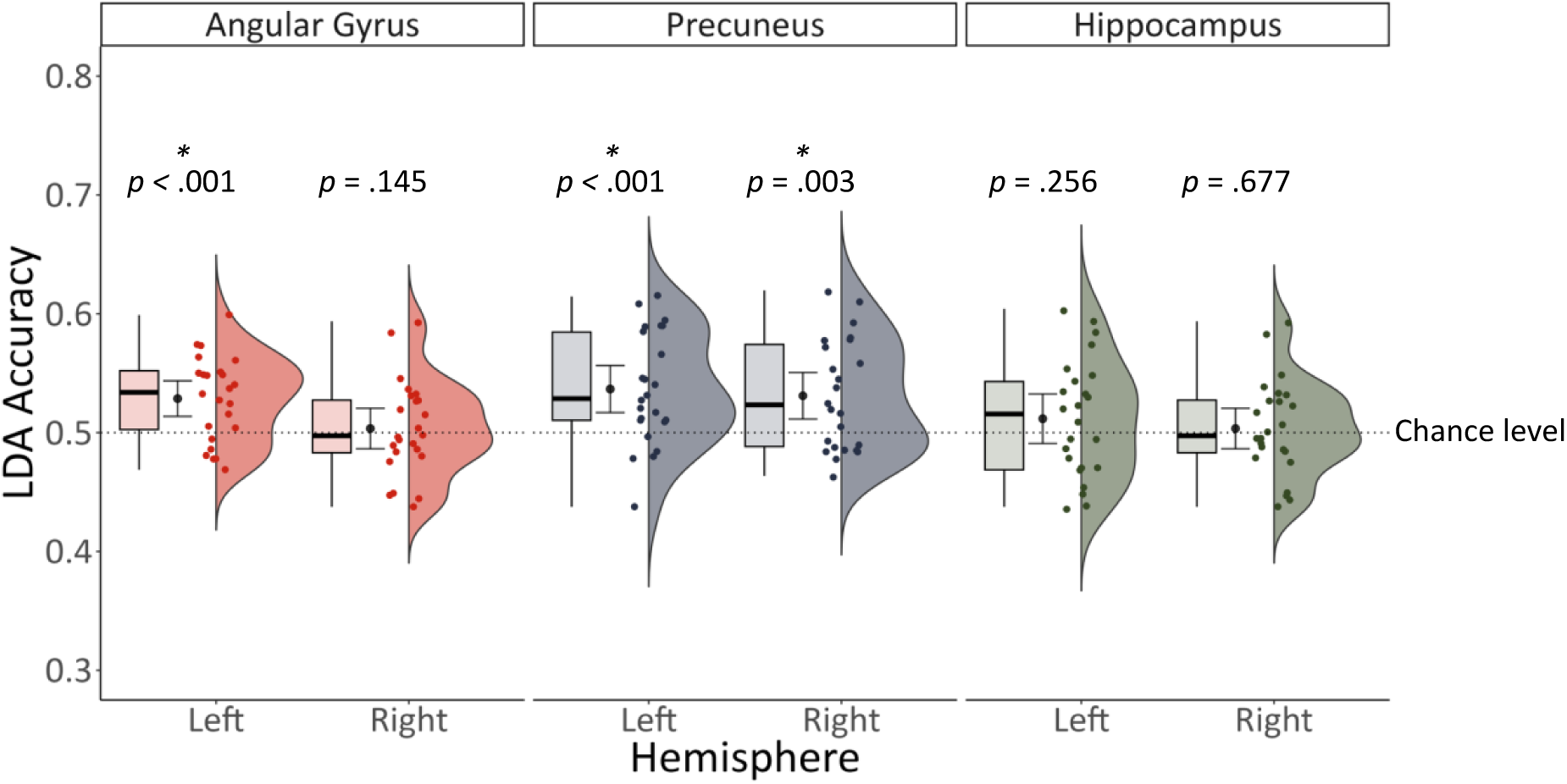
Decoding accuracy predicting embodiment was significantly greater than chance in the left angular gyrus, left precuneus, and right precuneus. 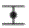 = mean decoding accuracy +/− 95% confidence interval.

#### Visual perspective x embodiment interaction

We investigated whether any of the six ROIs responded to the interaction between visual perspective and embodiment with two separate analyses. The first examined whether patterns of activity in each ROI predicted embodiment in the synchronous conditions only (i.e., sync in-body vs. sync out-of-body). We observed that the left angular gyrus was sensitive to this interaction effect, *t* (23) = 3.11, *p* = .005, Cohen’s d = 0.630, BF_10_ = 8.770 (*M* = .544, *SD* = .070; see Figure 7A). A Bayesian analysis indicated anecdotal evidence against greater than chance decoding accuracy in the right angular gyrus (BF_10_ = .932), and moderate evidence against greater than chance decoding accuracies in the precuneus (left: BF_10_ = .265, right: BF_10_ = .264) and hippocampus (left: BF_10_ = .256, right: BF_10_ = .225).

**Figure 7.**
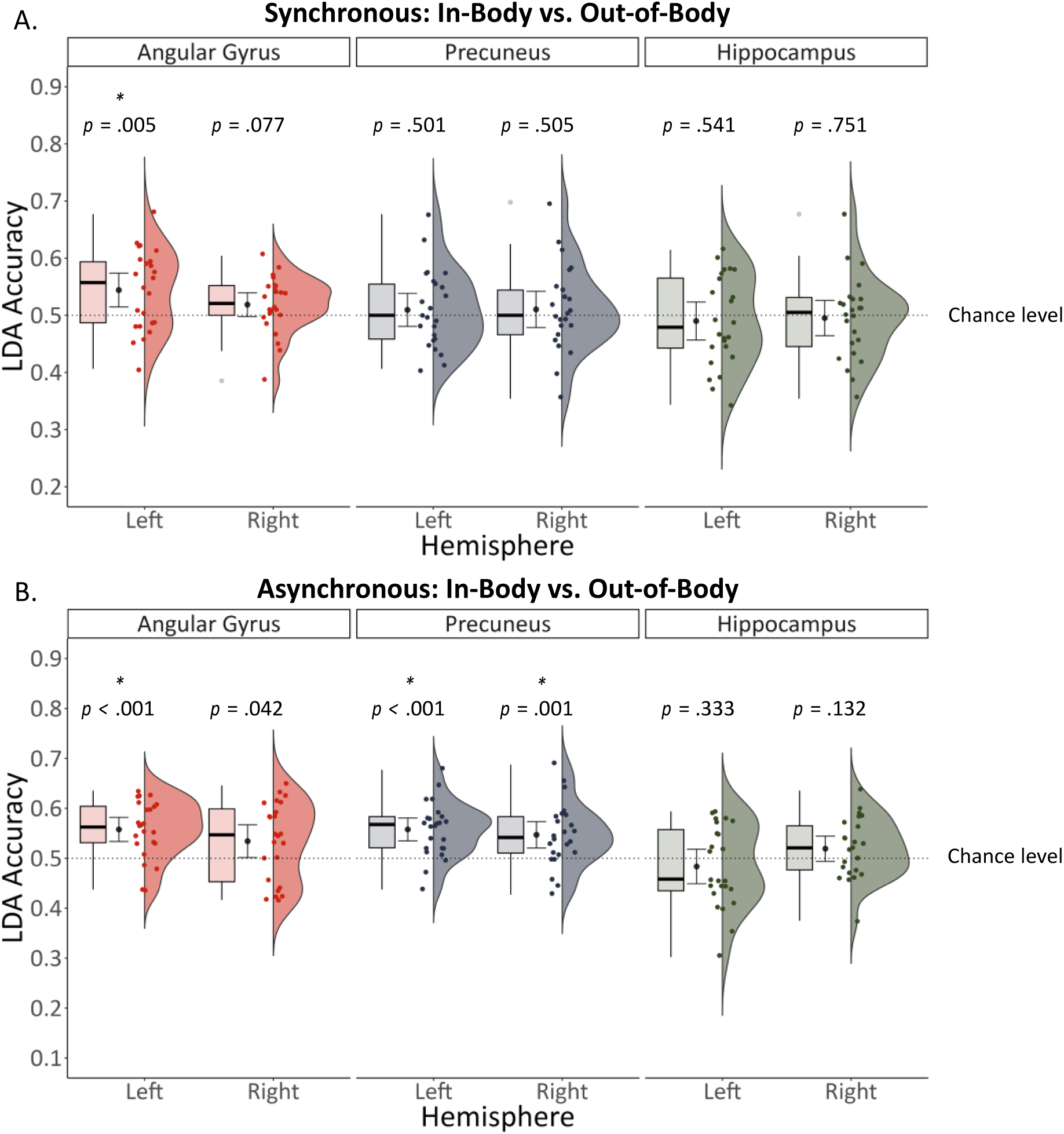
For synchronous conditions, patterns of activity during retrieval in the left angular gyrus predicted whether an event was associated with an in-body or out-of-body perspective. (A). For asynchronous conditions, patterns of activity in the left angular gyrus and bilateral precuneus predicted embodiment (B). 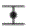 = mean decoding accuracy +/− 95% confidence interval.

The second investigated whether patterns of activity in each ROI predicted embodiment in the asynchronous conditions only (i.e., async in-body vs. async out-of-body). This effect was identified in the left angular gyrus *t*(23) = 4.964, *p* < .001, Cohen’s d = 1.018, BF_10_ = 495.841 (*M* = .558, *SD* = .057), left precuneus, *t*(23) = 5.221, *p* < .001, Cohen’s d = 1.074, BF_10_ = 880.160 (*M* = .558, *SD* = .054), and right precuneus, *t*(23) = 3.657, *p* = .001, Cohen’s d = .746, BF_10_ = 27.615 (*M* = .547, *SD* = .063) (see Figure 7B). A Bayesian analysis indicated weak evidence for above chance decoding accuracy in the right angular gyrus (BF_10_ = 1.507) and moderate evidence against above chance decoding accuracy in the hippocampus (right: BF_10_ = .632; left: BF_10_ = .333).

#### Whole-brain searchlight analyses

The whole brain searchlight analyses did not reveal any regions where patterns of activity could decode visual perspective, embodiment, or the interaction of the two after corrections for multiple comparisons (*p_FWE_* < .05). Due to the strict nature of the statistical thresholds required to correct for multiple comparisons across the entire brain, we applied a more liberal threshold of *p* <. 01 and cluster size > 5 voxels to descriptively investigate regions in each analysis that did not survive *FWE*-correction (see Supplemental Material).

## Discussion

The present study sought to understand how fundamental aspects of bodily selfhood, namely feelings of embodiment (i.e., self-location and body ownership) and visual perspective (i.e., in-body perspective, out-of-body perspective), experienced during the encoding of naturalistic events influence how that event is later represented in brain regions core to memory retrieval. Targeted region-of-interest multivariate analyses revealed that embodiment could be decoded from patterns of activity in the left angular gyrus and bilateral precuneus during retrieval. We further observed interactions between visual perspective and embodiment in these same regions. Visual perspective could be decoded from patterns of activity in the left angular gyrus for memories initially formed with both a strong and weak sense of embodiment. However, activity in the bilateral precuneus during retrieval predicted visual perspective for events associated with weak embodiment only.

While our behavioral manipulation of visual perspective was successful (i.e., memories encoded from a given perspective were retrieved from that same perspective), Bayesian analyses indicated only weak evidence for significant decoding of visual perspective alone, separated from embodiment, in the neuroimaging data. Further evidence is required to substantiate the involvement of the angular gyrus and precuneus in integrating visual perspective within memories of past events, isolated from embodiment. Our results do offer strong evidence that feelings of embodiment initially present during encoding are the predominant feature stored in posterior parietal memory traces for past events, and visual perspective is likely represented as a secondary, related component of bodily selfhood. The neuroimaging results are linked to our experimental manipulations of embodiment and visual perspective, rather than mediated by indirect influences in memory performance and subjective re-experiencing, as there were no initial differences between the conditions in memory accuracy, vividness, emotional intensity, or perceived memory accuracy. The visual perspective an event was encoded from determined the perspective at retrieval, which is consistent with previous literature (Bergouignan et al., 2022: Iriye & St. Jacques, 2021). Both immediately after encoding and during in-scanner retrieval, memories formed from an in-body perspective were retrieved from an in-body perspective, and memories formed from an out-of-body perspective were retrieved from an out-of-body perspective. Further, the interaction of visual perspective and embodiment during encoding influenced post-scanning memory accuracy. After repeated retrieval, cued recall accuracy was higher for memories encoded from an embodied in-body perspective, compared to an embodied out-of-body perspective. Together, the results reveal how the feeling of being located within and experiencing the world from the perspective of one’s own body determines the ability to accurately remember details of past events, the visual perspective they are re-experienced from at retrieval, and how past events are represented in brain regions essential for creating cohesive mental scenes based on egocentric reference frames.

We observed strong evidence that patterns of activity in the left angular gyrus reflected information pertaining to embodiment, and the interaction between embodiment and visual perspective. The present results align with theories that the angular gyrus integrates various sensory and spatial components of memories into a cohesive, egocentric framework, allowing individuals to subjectively relive these memories during retrieval (Bonnici et al., 2016, 2018; Ciaramelli et al., 2010b; Yazar et al., 2014, 2017). The angular gyrus, especially in the left hemisphere, is a “convergence zone” that combines several types of multimodal information (Seghier, 2023). Previous research has shown that the multisensory information integrated in this region includes visual and tactile signals related to bodily selfhood. For example, trial-by-trial fluctuations in the left angular gyrus activity predict the emergence of the feeling of illusory ownership over a limb (Chancel et al., 2022). Further, patterns of activity in the left angular gyrus present during the encoding of an immersive event reflect strong versus weak body ownership and are reinstated more during the retrieval of memories encoded with strong body ownership and high vividness (Iriye et al., 2024). Additionally, a meta-analysis found that bilateral angular gyri are implicated in mediating both feelings of bodily selfhood and episodic autobiographical memory (Bréchet et al., 2018). Thus, the results of the present study align with previous research identifying the angular gyrus as a hub of multisensory integration leading to feelings of embodiment within memories of past events.

At the same time, the present study offers novel insights into how the angular gyrus represents visual perspective. Specifically, we show that visual perspective is intertwined with feelings of embodiment in memories of past events such that evidence for above chance classifier performance was strongest when the visual perspective of a given memory was decoded considering the level of embodiment experienced during encoding. Earlier investigations that have manipulated either visual perspective or embodiment, but not both, miss a fundamental component of the information stored in this region. For example, earlier studies have suggested that the angular gyrus is crucial for creating a first-person vantage point within a mental scene, enabling subjective remembering during retrieval (Bonnici et al., 2016; 2018; Ciaramelli et al., 2010; Yazar et al., 2017). Specifically, applying continuous theta burst stimulation of the left angular gyrus decreases the likelihood of recalling memories from an in-body perspective, highlighting this region’s crucial role in forming a cohesive first-person perspective during retrieval. This account aligns with neuropsychological findings that report a correlation between disrupted angular gyrus activity and the occurrence of out-of-body experiences (Blanke et al., 2004; Blanke et al., 2002; Ionta et al., 2011), and that lesions here reduce first-person perspective imagery during spatial navigation (Ciaramelli et al., 2010). Based on previous research, the angular gyrus seems to play an essential role in forming a unified in-body perspective from which we perceive the world and re-experience memories. In contrast, the results of the present study suggest that the angular gyrus is involved in establishing in-body and out-of-body perspectives according to embodiment experienced during encoding (i.e., angular gyrus activity during retrieval predicted visual perspective for events encoded with both strong and weak feelings of embodiment separately). This finding suggests the angular gyrus is attuned to differences between in-body and out-of-body perspectives, and the sense of embodiment reactivated during retrieval. The angular gyrus’ ability to associate different visual perspectives with feelings of bodily selfhood may then enable one to flexibly shift between these viewpoints (St. Jacques et al., 2017). Thus, the current study enhances our understanding of egocentric representations in the angular gyrus by showing that this region instantiates both in-body and out-of-body visual perspectives and integrates them with feelings conveying embodiment while retrieving memories for past events. By explicitly manipulating visual perspective alongside feelings of embodiment in the present study, we reveal that information linked with visual perspective in memories is tightly linked with the feelings of body ownership and self-location initially experienced during encoding. These findings provide a more complete account of the role of the angular gyrus in memory for past events.

Like the angular gyrus, patterns of activity in the precuneus during memory retrieval also reflected embodiment at encoding, and the interaction between embodiment and visual perspective. However, unlike the angular gyrus, activity in the precuneus during memory retrieval reflected visual perspective only when the feeling of embodiment during initial encoding was weak. Direct electrical stimulation of the anterior precuneus bilaterally causes feelings of dissociation (Lyu and colleagues, 2023), suggesting that this region is sensitive to detachment from one’s body, which is required to adopt out-of-body perspectives in mental scenes. The precuneus is often associated with the ability to adopt a specific visual perspective in a memory or mental scene (Eich et al., 2009b; Freton et al., 2014b; Grol et al., 2017; Iriye & St. Jacques, 2020) and elaborate upon its visual details (Daselaar et al., 2008; Fuentemilla et al., 2014; Gardini et al., 2006; Söderlund et al., 2012). However, we observed that information related to visual perspective could only be reliably detected when also including information connected to embodiment at encoding. Bayesian analyses revealed that we could not conclusively determine whether visual perspective on its own, separated from embodiment, could be decoded from precuneus activity.

This finding implies that previous studies on the role of visual perspective in memory may have also inadvertently been probing associated changes in embodiment (i.e., self-location, body ownership) that come with adopting a given visual perspective in memory.

Moreover, previous results of the involvement of the precuneus during the instantiation of visual perspective in mental scenes can be accounted for by discrepant demands required to shift between visual perspectives (St. Jacques et al., 2017; 2018) and manipulate mental scenes inside an egocentric reference frame (Byrne et al., 2007). In support of this theory, St. Jacques, Szupunar, and Schacter (2017) showed that the posterior parietal cortex, including the precuneus and angular gyrus, is engaged when participants actively change their visual perspective from an in-body perspective to an observer perspective during memory retrieval. Additionally, they observed that the level of precuneus involvement in this perspective shift predicted decreases in emotional intensity at retrieval and ensuing changes in the main visual perspective of autobiographical memories. This finding has been replicated by additional research, which demonstrated that a shift in visual perspective increased activation of the precuneus, irrespective of the direction of the perspective shift (St. Jacques et al., 2018). These findings align with theories of memory and imagination, which propose that egocentric frameworks formed during long term memory retrieval in the precuneus can be modified when individuals envision the potential available movements within a mental scene (Byrne et al., 2007). These processes might be more actively engaged when individuals adopt an out-of-body perspective during memory retrieval, as retrieving memories from a new self-location often necessitates the updating of internal representations of the mental scene. The present study did not incur greater demands upon mental transformation processes while retrieving memories from an out-of-body perspective as the events were initially formed from an out-of-body perspective.

Consequently, participants did not need to adjust their mental scene to accommodate an updated perspective. Here, the lack of strong evidence supporting significant decoding of visual perspective on its own separated from embodiment may be attributable to the sensitivity of this region to mental transformation processes of mental scenes, which have been different between visual perspectives in earlier investigations (Grol et al., 2017; Iriye & St. Jacques, 2020; St. Jacques et al., 2017; 2018). Directly manipulating mental transformation processes needed to establish new visual perspectives in mental scenes should be incorporated into future research to clarify the role of the precuneus in memory retrieval.

Turning to the hippocampus, we did not find activity in this region predicted visual perspective, embodiment, or their interaction during memory retrieval. Previous research has shown that retrieving memories of events encoded from an out-of-body perspective leads to delayed activation of the left posterior hippocampus, linked to decreased vividness and poorer recall of episodic details compared to memories encoded from an in-body perspective (Bergouignan et al., 2014). Further, bilateral hippocampal activity during encoding distinguishes between memories being formed with strong versus weak body ownership, which is then reactivated to a greater degree during the retrieval of memories formed with strong body ownership and high vividness (Iriye et al., 2024). Conflicting results between these two previous studies and the present study may be due to reduced durability of memory traces for events encoded from an out-of-body perspective.

Bergouignan and colleagues (2014) found that memory accuracy and vividness were lower for events encoded from an out-of-body relative to in-body perspective. In the present study, immediate memory performance and subjective re-experiencing was equated between conditions by repeating the word games several times and asking participants to retrieve mental images of each game immediately after encoding. Iriye and St. Jacques (2021) found that the durability of out-of-body perspective ratings decreased over the course of a week, further suggesting that memory traces for out-of-body events are less stable over time. Iriye and colleagues (2024) manipulated body ownership, but not visual perspective, and did not observe differences in memory accuracy between conditions.

Together, these studies suggest that out-of-body perspectives may lead to less durable memory traces, which may contribute to changes in the involvement of the hippocampus and may have also led to inconclusive results regarding the decoding of visual perspective in the angular gyrus and precuneus. Regardless, the results of the present study underscore that bodily selfhood is integrated within memories of past events in regions outside of the medial temporal lobes and in posterior parietal cortex.

Our results generate original insights into clinical reports of disorders exhibiting both abnormal body experiences and memory impairments. For example, disruptions in own-body perception are a core symptom of schizophrenia (Costantini et al., 2020; Schultze-Lutter, 2009), which can involve a weakened sense of body ownership and blurred boundary between self and the external world (Di Cosmo et al., 2018; Thakkar et al., 2011; van der Weiden et al., 2015). At the same time, schizophrenia is often associated with impairments in episodic memory, which have been linked to an inability to bind object level information with spatial contextual information during encoding (Aleman et al., 1999; Brébion et al., 1997; Rushe et al., 1999; Talamini et al., 2010). Deficits in remembering verbally presented information, such as the word games in the present study, during childhood are also a key predictor of the development of schizophrenia (Erlenmeyer-Kimling et al., 2000). The paradigm of the present study allows for the induction of abnormal body experiences (i.e., weakened body ownership, shifts in self-location, and out-of-body perspectives) in healthy individuals. Using this innovative paradigm that creates highly realistic, immersive memories tightly controlled in a lab setting, we show that encoding events from an out-of-body perspective leads to reduced recall of event details compared to when events are encoded from a perspective within one’s body, which mimics reported impairments in episodic memory in patients with schizophrenia. Previous models of impaired binding processes during memory encoding in schizophrenia have focused on connectivity within medial temporal lobe regions (Talamini et al., 2005, 2010; Talamini & Meeter, 2009). However, our results indicate that posterior parietal regions including the angular gyrus and precuneus contain information related to embodiment and visual perspective that are critical to building egocentrically defined reference frames that facilitate memory retrieval and should be considered in medio-temporal lobe models of schizophrenia. While the present study offers a novel paradigm to mimic the disturbances in body ownership and memory impairments commonly observed in schizophrenia, which provides a promising avenue to investigate the mechanism of this interaction, it focuses on healthy participants with a relatively small sample size. Future research could expand on this paradigm in clinical populations to further clarify how visual perspective and embodiment interact to support memory of past events.

Our behavioural results indicated that differences in memory related to changes in bodily selfhood during memory formation may emerge over time. The cued recall test in the post-scanning session revealed that memory accuracy was higher for in-body compared to out-of-body perspectives for memories formed with strong embodiment. This finding is consistent with previous reports that disrupting feelings of bodily selfhood is linked to deficits in episodic memory (Bergouignan et al., 2014; Bréchet et al., 2018, 2019; Iriye et al., 2022). For example, Bergouignan and colleagues (2014) also show that encoding events from an embodied third-person perspective is linked to reduced episodic remembering, compared to an embodied first-person perspective. In contrast, we did not observe any differences between conditions in terms of memory vividness, emotional intensity, or belief in memory accuracy either immediately after encoding or after repeated retrieval. Previous research has shown higher memory vividness, emotional intensity, and belief in memory accuracy for memories encoded with strong versus weak embodiment, but only with a one-week delay between encoding and retrieval (Iriye & Ehrsson, 2022). Encoding and retrieval took place on the same day in the present study, suggesting that a delay is required for the effects of embodiment on memory phenomenology to manifest. Our results are also in line with prior evidence demonstrating that encoding events from an in-body versus out-of-body perspective does not influence vividness, emotional intensity or reliving either immediately after encoding or after a one-week delay (Iriye & St. Jacques, 2021). Further, the present study included a manipulation of visual perspective that was not part of the paradigm in Iriye & Ehrsson (2022), which may have led to differential effects on memory phenomenology. The present study is the first to simultaneously manipulate visual perspective and embodiment to examine influences memory properties.

### Limitations

The current results should be interpreted considering the study’s limitations. We aimed to create events that were as close to real life scenarios as possible while retaining the ability to reliably experimentally manipulate visual perspective and embodiment. This meant that participants were required to remain still during the word games since movement would have influenced feelings of agency over one’s body, thereby disrupting the body illusions (Tsakiris, Prabhu, & Haggard, 2006). The participants’ inability to move during the word games limits the naturalistic experience of the experiment.

However, the dialog in real-time between the experimenter and participant provided a realistic, dynamic social interaction while the association of each word game to a unique object within the cabinet made the games visually lifelike as well as memorable. Future experiments adopting advanced virtual technologies will enhance the ability of experimenters to manipulate complex cognitive phenomena within increasingly naturalistic environments.

A second potential limitation is that the immediate cued recall test and subjective ratings may have influenced memory retrieval in-scanner and the post-scanning cued recall test and subjective ratings, which limited our ability to capture potential differences in memory performance. The multiple retrievals were necessary to ensure that memory accuracy and subjective components of remembering were equivalent across the conditions immediately after encoding to isolate the effects of our bodily selfhood manipulation on neural representations during memory retrieval. Further, repeated retrievals of each event during scanning were necessary to obtain sufficient power to detect effects of visual perspective and embodiment on remembering. Multiple retrieval repetitions can lead to repetition suppression effects. However, since the decoding analyses were designed to identify unique patterns of brain activity associated with each condition, and not repetition suppression effects, repeated retrieval likely did not affect the fMRI results. Although repeated retrieval may have led to re-encoding of some memories—particularly for less stable out-of-body events—we consider it unlikely that this confound fully accounts for the observed decoding effects. Our multivariate classification approach was designed to identify condition-specific patterns of neural activity, and cross-validation ensured that classifier performance was not driven by trial-level fluctuations. Further, decoding analyses are sensitive to representational differences rather than signal amplitude and, thus, are not directly confounded by repetition suppression or enhancement effects. While re-encoding may have occurred, it would not explain the condition-specific neural patterns unless it introduced consistent structure across trials within a condition. Future research examining the effects of bodily selfhood manipulations in the absence of rehearsal would be of interest to better understand how perspective and embodiment contribute to changes in memory performance.

Finally, another limitation is that we did not observe any regions that could predict visual perspective, embodiment, or their interaction during memory retrieval at the whole-brain level, which may be due to our limited sample size. However, whole-brain searchlight analyses involve conservative corrections for multiple comparisons for a high number of statistical tests performed over the entire brain, which may have led to false negative findings (Etzel, Zacks, & Braver, 2013). This idea is confirmed by our observations that applying a more liberal statistical threshold implicated the left angular gyrus and left precuneus in decoding visual perspective, and the left angular gyrus and right precuneus in decoding embodiment (see Supplemental Material). We used the ROI approach to specifically test our hypotheses about the angular gyrus, precuneus, and hippocampus based on existing literature to limit the number of required statistical tests and limit the influence of the multiple corrections problem. We chose to investigate lateralized contributions of each ROI to visual perspective and embodiment by separating each ROI into a left and right hemisphere motivated by prior literature suggesting hemispheric asymmetries in these processes. However, decoding based on bilateral ROI patterns is an important direction for future work that could reveal integrative conributions that are not detectable through separate hemispheric analyses.

### Conclusion

In sum, the present study bridges the typically disconnected areas of memory and embodiment by offering fresh insights into how a person’s distinct perspective on the world and feeling of bodily selfhood together shape neural activity patterns involved in retrieval. Patterns of activity in the left angular gyrus and bilateral precuneus during the retrieval of lifelike events predicted feelings of embodiment experienced during encoding. While the left angular gyrus was involved in the retrieval of events from in-body and out-of-body perspectives regardless of the strength of the embodiment at encoding, neural activity in the bilateral precuneus differentiated visual perspectives only for memories with weak embodiment. Ultimately, this line of research elucidates how an elusive, multifaceted sense of bodily self becomes incorporated within memories of the personal past to structure higher levels of selfhood contingent on autobiographical memories. Here, we highlight brain regions that may be implicated in mediating episodic memory impairments reported in disorders involving depersonalization and dissociation, such as schizophrenia (Borda & Sass, 2015; Prebble et al., 2013; Sierra & David, 2011).

## Data Code and Availability

Original data and code are available upon reasonable request through contact with the corresponding author.

## Author Contributions

Both Heather Iriye and Peggy L. St. Jacques participated in conceptualization of the experiment and the design of its methodology. Heather Iriye collected the data, conducted the formal analysis, visualized the results, and wrote the original draft of the manuscript. Peggy L. St Jacques reviewed and edited the manuscript, performed supervision, was the project administrator, and acquired the funding.

## Acknowledgments

We thank radiographers Janice Bush, James Hunter, and Marta Andrade, at the Clinical Imaging Science Centre, University of Sussex, for their assistance with fMRI data collection.

