## Supplemental Table 1 for "An embodied perspective: angular gyrus and precuneus decode selfhood in memories of naturalistic events"

### Supplemental Material

*Supplemental Table 1: List of Statements Used in the Truth Game*

| Memory Cue | Statement 1 | Statement 2 | Statement 3 |
|---|---|---|---|
| *Groups* | A group of crows is called a murder. | A group of owls is called a parliament. | A group of bats is called a bevvy. (*False)* |
| *Brighton* | The Brighton Sea Life Centre is the world's oldest aquarium. | The University of Sussex was Established in 1961. | Brighton is the 2nd happiest place to live in the UK. (*False)* |
| *Berries* | A banana is a berry. | An avocado is a berry. | A strawberry is a berry. (*False)* |
| *Favorites* | My favorite sport is baseball. | My favorite beverage is coffee. | My favorite colour is purple. (*False)* |

*Supplemental Table 2: Cued Recall Questions*

| Game | Encoding Session Question | Post-Scanning Session Question |
|---|---|---|
| *I Spy* | What was your first guess? | What was your last guess? |
| *Categories* | What was the first example you came up with? | What was the last example you came up with? |
| *Sentence Construction* | What was the first word of the sentence? | What was the last word of the sentence? |
| *Truth Game* | What was the first statement? | What was the last statement? |

*Supplemental Table 3: Memory Test Results Not Reported in Main Text*

| Session | Measure | Condition | Mean (s.d.) |
|---|---|---|---|
| *Immediate Memory Test* | *Memory Accuracy* | In-Body Sync | .98 (.07) |
| | | In-Body Async | .96 (.10) |
| | | Out-of-Body Sync | .97 (.08) |
| | | Out-of-Body Async | .99 (.05) |
| | *1PP* | In-Body Sync | 5.98 (1.16) |
| | | In-Body Async | 5.75 (1.37) |
| | | Out-of-Body Sync | 3.91 (1.27) |
| | | Out-of-Body Async | 3.56 (1.71) |
| | *3PP* | In-Body Sync | 1.63 (1.10) |
| | | In-Body Async | 1.87 (1.09) |
| | | Out-of-Body Sync | 4.27 (1.52) |
| | | Out-of-Body Async | 3.75 (1.78) |
| | *Vividness* | In-Body Sync | 5.40 (.86) |
| | | In-Body Async | 5.32 (.99) |
| | | Out-of-Body Sync | 5.37 (.96) |

| | | | |
|---|---|---|---|
| | | Out-of-Body Async | 5.24 (.92) |
| | *Emotional Intensity* | In-Body Sync | 3.34 (1.44) |
| | | In-Body Async | 3.45 (1.62) |
| | | Out-of-Body Sync | 3.29 (1.43) |
| | | Out-of-Body Async | 3.12 (1.35) |
| | *Belief in Memory* | In-Body Sync | 5.41 (.79) |
| | *Accuracy* | In-Body Async | 5.52 (1.06) |
| | | Out-of-Body Sync | 5.12 (1.05) |
| | | Out-of-Body Async | 5.40 (.88) |
| *In-Scanner* | *1PP* | In-Body Sync | 2.99 (.60) |
| | | In-Body Async | 3.06 (.60) |
| | | Out-of-Body Sync | 2.45 (.71) |
| | | Out-of-Body Async | 2.51 (.67) |
| | *3PP* | In-Body Sync | 1.70 (.42) |
| | | In-Body Async | 1.65 (.41) |
| | | Out-of-Body Sync | 2.25 (.68) |
| | | Out-of-Body Async | 2.19 (.69) |
| | *Vividness* | In-Body Sync | 3.07 (.48) |
| | | In-Body Async | 3.08 (.55) |
| | | Out-of-Body Sync | 2.97 (.54) |
| | | Out-of-Body Async | 3.05 (.43) |
| *Post-Scanning* | *Emotional Intensity* | In-Body Sync | 3.33 (1.38) |
| | | In-Body Async | 3.39 (1.45) |
| | | Out-of-Body Sync | 3.26 (1.42) |
| | | Out-of-Body Async | 3.26 (1.43) |
| | *Belief in Memory* | In-Body Sync | 5.35 (.88) |
| | *Accuracy* | In-Body Async | 5.32 (.91) |
| | | Out-of-Body Sync | 5.04 (1.29) |
| | | Out-of-Body Async | 5.27 (.91) |

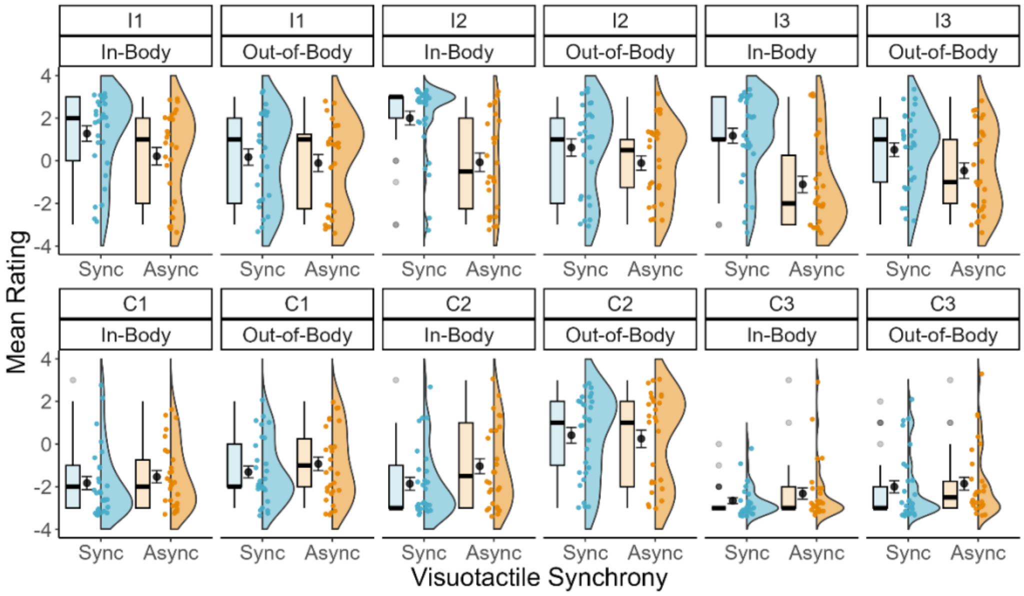


*Supplemental Figure 1*. Individual statement ratings for the illusion induction questionnaire.

### Supplemental Results

Here we report results after removing outliers from the analyses. Including and excluding outliers led to the same pattern of results for each analysis and the conclusions remain the same. Whole brain searchlight results at the liberal threshold of $p$ <. 01, cluster size ($K$) > 5 voxels are also reported for descriptive purposes.

#### Behavioral Results

**Event encoding: Illusion Induction Questionnaire:** Three outliers were removed from the data. A 2 (Embodiment: sync, async) x 2 (Visual Perspective: in-body, out-of-body) repeated measures ANOVA on average illusion minus control statement ratings found main effects of embodiment, $F(1, 20) = 29.29$, $p < .001$, $\eta_p 2 = .59$, $BF_{10} = 370.20$ , and visual perspective, $F(1, 20) = 29.60$, $p < .001$, $\eta_p 2 = .60$, $BF_{10} = 34. 92$. The interaction was marginally significant, $F (1, 20) = 4.15$, $p = .06$, $\eta_p 2 = .17$ (although a Bayesian analysis indicates very strong evidence against the null hypothesis, $BF_{10} = 36603.35$). Synchronous compared to asynchronous visuotactile stimulation was linked to greater illusion minus control ratings, collapsed across visual perspectives, $t (20) = 5.11$, $p < .001$, Cohen's d = 1.09, $BF_{10} = 9610.05$, demonstrating that the illusion worked as expected in both in-body and out-of-body conditions. In-body compared to out-of-body perspectives led to higher illusion minus control ratings, $t (20) = 4.52$, $p < .001$, Cohen's d = .75, $BF_{10} = 262.68$.

**Immediate memory test**. Eight outliers were removed from the data. A 2 x 2 repeated measures ANOVA with visual perspective (in-body, out-of-body) and embodiment (sync, async) conducted on average cued recall accuracy scores did not reveal a main effect of visual perspective, $F(1,15) = 2.14$, $p = .16$, $\eta_p 2 = .13$, embodiment, $F(1,15) < .001$, $p = 1.00$, $\eta_p 2 < 0.001$, or an interaction, $F(1,15) < .001$, $p = 1.00$, $\eta_p 2 < .001$. A Bayesian analysis indicated anecdotal evidence for the null hypothesis for visual perspective ($BF_{10}$ = .63) and embodiment ($BF_{10}$ = .33), and strong evidence for the null hypothesis for the interaction term ($BF_{10}$ = .07). Thus, immediate memory recall was equal between conditions.

To investigate whether the visual perspective participants adopted at immediate retrieval differed according to condition, we conducted a 2 (visual perspective: in-body, out-of-body) x 2 (embodiment: sync, async) repeated measures ANOVA for 1PP and 3PP ratings separately. Three outliers were removed from the data. There was a significant main effect of visual perspective, $F$ (1,20) = 72.03, $p$ < .001, $\eta_p 2$ = .78, $BF_{10}$ = 404 586.66, where 1PP ratings were higher in the in-body ($M$ = 6.13, $SD$ = 1.07) compared to out-of-body condition ($M$ = 3.80, $SD$ = 1.65 ). There was no main effect of embodiment, $F$(1,20) = 1.72, $p$ = .21, $\eta_p 2$ = .08, $BF_{10}$ = .60, or interaction, $F$(1,20) = .01, $p$ = .91, $\eta_p 2$ < .001, $BF_{10}$ = .17 (although a Bayesian analysis indicated very strong evidence that a model including the interaction term was better at explaining the data than the null model, $BF_{10}$= 61 756.91). For the 3PP ratings, two outliers were removed from the data. There was a significant main effect of visual perspective, $F$ (1,21) = 74.51, $p$ < .001, $\eta_p 2$ = .78, $BF_{10}$ = 761 436.06, where 3PP ratings were higher in the out-of-body ($M$ = 3.88, $SD$ = 1.86) compared to in-body condition ($M$ = 1.49, $SD$ = 0.84 ). There was no main effect of embodiment, $F$(1,21) = 0.26, $p$ = .62, $\eta_p 2$ = .01, $BF_{10}$ = .30, or interaction, $F$(1,21) = 4.14, $p$ = .06, $\eta_p 2$ = .17, $BF_{10}$ = .56 (although a Bayesian analysis indicated very strong evidence that a model including the interaction term was better at explaining the data than the null model, $BF_{10}$ = 403 694.53).

For the vividness ratings, one outlier was removed from the data. A 2 (visual perspective: in-body, out-of-body) x 2 (embodiment: sync, async) did not find a main effect of visual perspective, $F$(1,22) = .80, $p$ = .27, $\eta_p 2$ = .04, $BF_{10}$ = .49, or visuotactile stimulation,

*F*(1,22) = 1.83, *p* = .19, $\eta_p2$ = .08, $BF_{10}$ = .30, and no interaction, *F*(1,22) = .30, *p* = .590, $\eta_p2$ = .01, $BF_{10}$ = .02.

**In-scanner subjective ratings.** The analysis of in-scanner visual perspective ratings revealed the same pattern of effects observed in the immediate memory test. One outlier was removed from the 1PP in-scanner ratings. There was a main effect of visual perspective, *F* (1,22) = 27.93, *p* < .001, $\eta_p2$ = .56, $BF_{10}$ = 541 129.32, indicating higher ratings in the in-body (*M* = 3.12, *SD* = 0.91) compared to out-of-body condition (*M* = 2.53, *SD* = 1.06). There was no main effect of embodiment, *F*(1,22) = .31, *p* = .31, $\eta_p2$ =.04, $BF_{10}$ = .60, or interaction, *F*(1,22) < .001, *p* =.99, $\eta_p2$ <.001 (although a Bayesian analysis indicated very strong evidence that a model including the interaction term was better at explaining the data compared to the null model, $BF_{10}$ = 94 768.04). Two outliers were removed from the 3PP in-scanner ratings. There was a main effect of visual perspective, *F* (1,21) = 5.18, *p* < .001, $\eta_p2$ =.52, $BF_{10}$ = 29 694.73, indicating higher ratings in the in-body (*M* = 2.13 , *SD* = 1.08) compared to out-of-body condition (*M* = 1.63, *SD* = .87). There was no main effect of embodiment, *F*(1,21) = .55, *p* = .47, $\eta_p2$ =.03, $BF_{10}$ = 1.12, or interaction, *F*(1,21) = .40, *p* =.54, $\eta_p2$ =.02 (although a Bayesian analysis indicated very strong evidence that a model including the interaction term was better at explaining the data compared to the null model, $BF_{10}$ = 11 929.62). Turning to the in-scanner vividness ratings, one outlier was removed from the data. There was no main effect of visual perspective, *F* (1,22) = 3.90, *p* = .06, $\eta_p2$ = .15, $BF_{10}$ = .85, embodiment, *F* (1,22) = .71, *p* = .41, $\eta_p2$ = .03, $BF_{10}$ = .40, or interaction, *F* (1,22) = .24, *p* = .63, $\eta_p2$ = .01, $BF_{10}$ = .12.

**Post-scanning cued recall and subjective ratings.** Three outliers were removed from the analysis of post-scanning cued recall accuracy. There was a marginally significant effect of visual perspective, *F* (1,20) = 4.03, *p* = .06, $\eta_p2$ = .17, $BF_{10}$ = 1.30. There was a significant interaction between visual perspective and embodiment, $F$ (1,20) = 5.39, $p$ = .03, $\eta_p 2$ = .21, $BF_{10}$ = 2.28. The in-body sync condition ($M$ = .99, $SD$ = .06) was associated with higher cued recall accuracy compared to the out-of-body sync condition ($M$ = .89, $SD$ = .15), $p$ = .046, $BF_{10}$ = 6.27. Turning to the belief in memory accuracy ratings, one outlier was removed from the analysis. There were no main effects of visual perspective, $F$ (1,22) = 1.02, $p$ = .32, $\eta_p 2$ = .04, $BF_{10}$ = .43, embodiment, $F$(1,22) = .31, $p$ = .59, $\eta_p 2$ = .01, $BF_{10}$ = .34, or interaction, $F$(1,22) = .69, $p$ = .41, $\eta_p 2$ = .03, $BF_{10}$ = .06.

**Multivariate Decoding Analyses Results: ROI Analyses**

**Visual perspective.** Two outliers were removed from the data of the right precuneus, but did not affect the results, $p$ = .026, $BF_{10}$ =2.28 (i.e., patterns of activity did not differentiate between visual perspectives after correcting for multiple comparisons, but the Bayesian analysis provided weak evidence for above chance decoding accuracy in this region).

**Visual perspective x embodiment interaction.** One outlier each in the right angular gyrus, right precuneus, and right hippocampus was removed from the analysis decoding in-body versus out-of-body perspectives in the synchronous conditions. Removing these outliers did not affect the pattern of results (right angular gyrus: $p$ = .02, $BF_{10}$ = 2.98; right precuneus: $p$ = .77, $BF_{10}$ = .24; right hippocampus: $p$ = .28, $BF_{10}$ = .53). Namely, none of these ROIs exhibited above chance decoding accuracy after correcting for multiple comparisons.

**Games.** Significant decoding of game type was observed in bilateral angular gyrus (right: $M$ = .387, $SD$ = .105, $t$(23) = 6.36, $p$ < .001, Cohen's d = 1.298 , $BF_{10}$ = 10839.283; left: $M$ = .411, $SD$ = .110, $t$(23) = 7.168, $p$ < .001 , Cohen's d = 1.463, $BF_{10}$ = 60469.728), bilateral precuneus (right: $M$ = .397, $SD$ = .117, $t$(23) = 6.129, $p$ < .001 , Cohen's d = 1.251, $BF_{10}$ =

6565.225; left: $M$ = .432, $SD$ = .121, $t(23)$ = 7.336, $p < .001$ , Cohen's d = 1.497 , $BF_{10}$ = 85736.095), and left hippocampus ($M$ = .269, $SD$ = .031 , $t(23)$ = 2.909, $p$ = .008, Cohen's d = .594, $BF_{10}$ = 5.897) (see Supplementary Figure 2). For these ROIs where classifier performance was significant, we compared classifier accuracy scores between game types to assess potential differences in classifier performance, which may have influenced the results of the main decoding analyses. The main effect of game was significant in the right angular gyrus ($F$ (3,69) = 2.94, $p$ = .039, $\eta_p 2$ = .113, $BF_{10}$ = 1.297). However, follow-up t-tests corrected for multiple comparisons with the Bonferroni-Holm method did not reveal any significant differences in classifier performance between games (all $p$'s > .081), which is in line with the Bayes factor of the main effect indicating only anecdotal evidence for a difference. Classifier accuracy was equal across game types in the left angular gyrus ($F(3,69)$ = .456 , $p$ = .714, $\eta_p 2$ = .019, $BF_{10}$ = .092), bilateral precuneus (right: $F(3,69)$ = 1.835, $p$ = .149, $\eta_p 2$ = .074, $BF_{10}$ = .405; left: $F(3,69)$ = 1.284 , $p$ = .287, $\eta_p 2$ = .053, $BF_{10}$ = .223), and the left hippocampus ($F(3,69)$ = 1.772, $p$ = .161, $\eta_p 2$ = .072, $BF_{10}$ = .472). See Supplementary Figure 3 for confusion matrices displaying classification accuracies according to game type for each ROI. Thus, differences in visual perspective between game types were unlikely to influence the results of the main ROI analyses decoding visual perspective and embodiment.

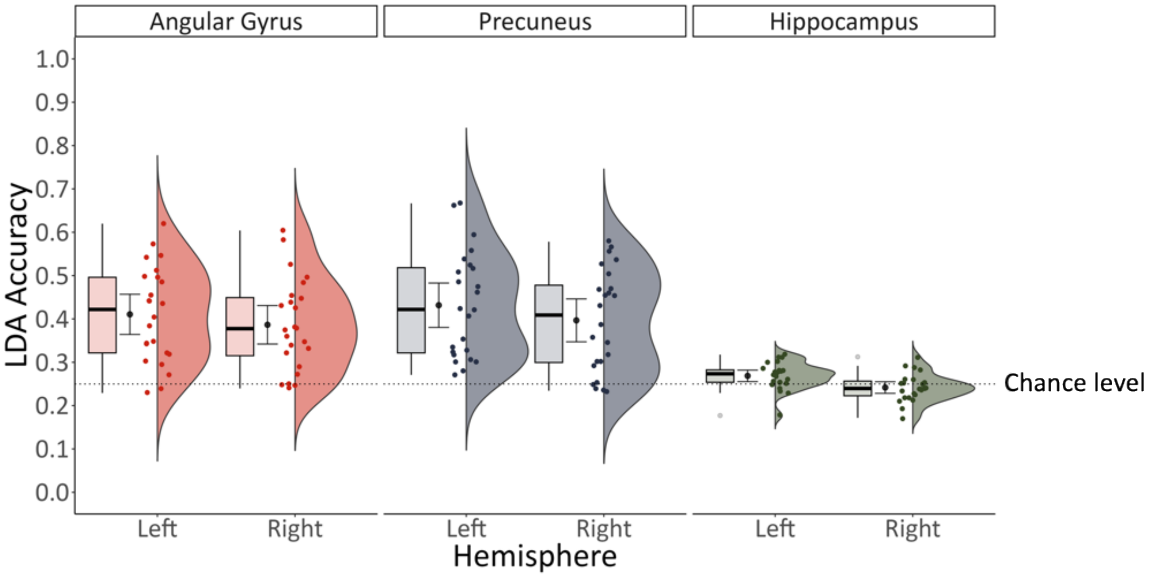


*Supplementary Figure 2*. Game type was predicted from patterns of activity in bilateral precuneus, bilateral angular gyrus, and the left hippocampus.

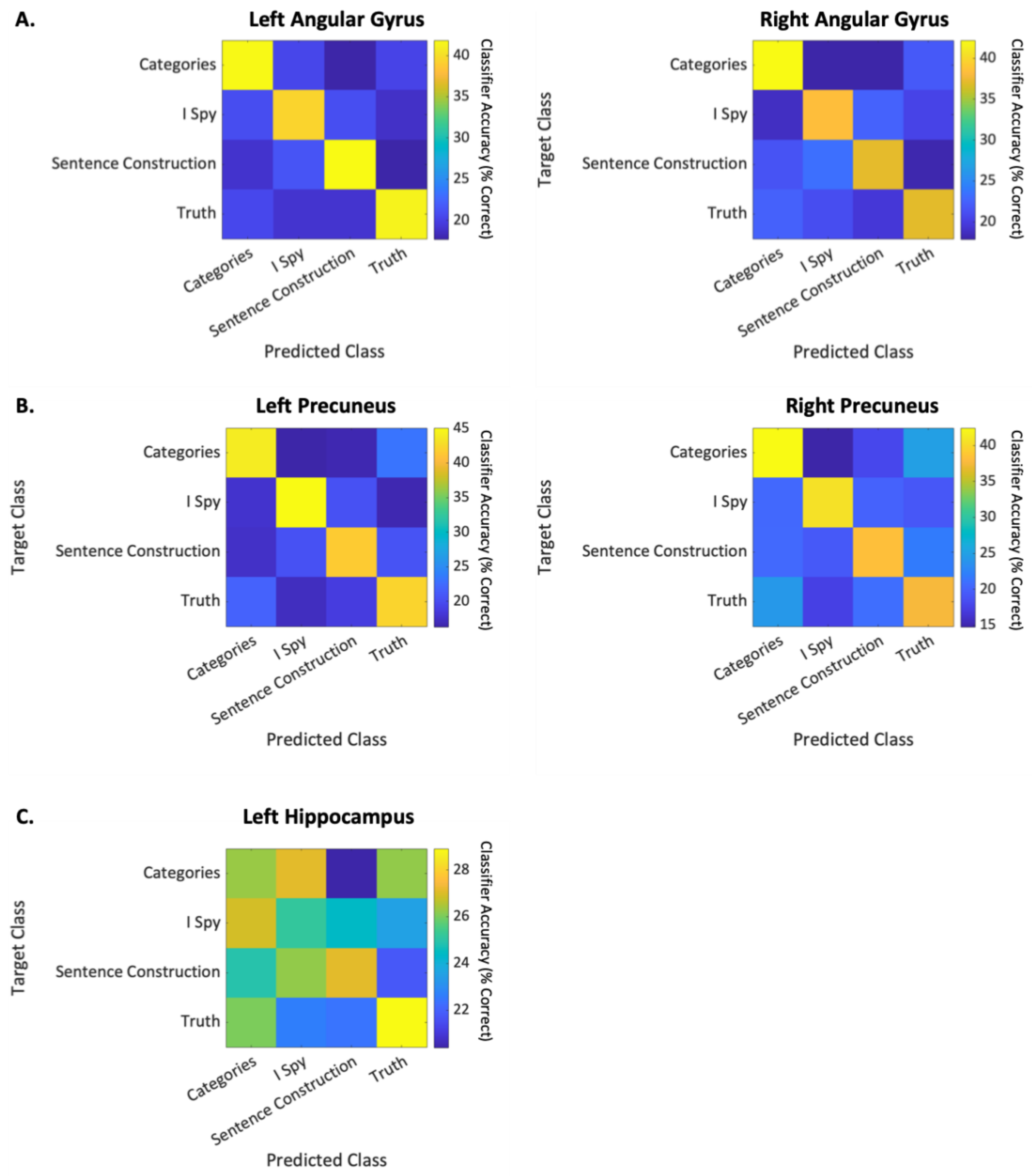


*Supplementary Figure 3*. Classification accuracies across game types were equal in the angular gyrus (A), precuneus (B), and hippocampus (C).

### Multivariate Decoding Analysis Results: Whole Brain Analysis

**Visual perspective.** Due to the conservative nature of the whole brain searchlight approach, we lowered the statistical threshold to $p < .01$ and $K > 5$ voxels to investigate regions that may be implicated in visual perspective despite not surviving corrections for multiple comparisons across the entire brain. In line with our predictions and ROI results reported in the main text, activity in the left angular gyrus and left precuneus predicted a memory's visual perspective during retrieval (see Supplemental Figure 4 & Supplemental Table 4). The left fusiform gyrus, left precentral gyrus, and left frontal regions were also identified.

*Supplemental Table 4: Whole Brain Searchlight Analysis, Visual Perspective*

| Region | BA | K | $p_{uncorr}$ | T | X | Y | Z |
|---|---|---|---|---|---|---|---|
| Angular Gyrus | 39 | 5 | < .001 | 5.33 | -48 | -68 | 32 |
| Fusiform Gyrus | 37 | 5 | <.001 | 5.15 | -44 | -52 | -18 |
| Precentral Gyrus | 4 | 11 | <.001 | 4.93 | -60 | 0 | 20 |
| Superior Frontal Gyrus | 6 | 5 | <.001 | 4.86 | -16 | 18 | 66 |
| Inferior Frontal Gyrus (Pars Triangularis) | 45 | 12 | <.001 | 3.93 | -50 | 26 | 12 |
| Inferior Frontal Gyrus (Pars Triangularis) | 45 | | .001 | 3.48 | -44 | 20 | 16 |
| Middle Frontal Gyrus | 10 | 6 | .001 | 3.72 | -32 | 58 | 22 |
| Precuneus | 7 | 6 | .001 | 3.09 | -2 | -66 | 26 |
| Middle Frontal Gyrus | 9 | 5 | .003 | 2.70 | 46 | 30 | 28 |

* The statistical map was thresholded at $p < .01$, $K > 5$ for display purposes. No clusters survived FWE-correction.

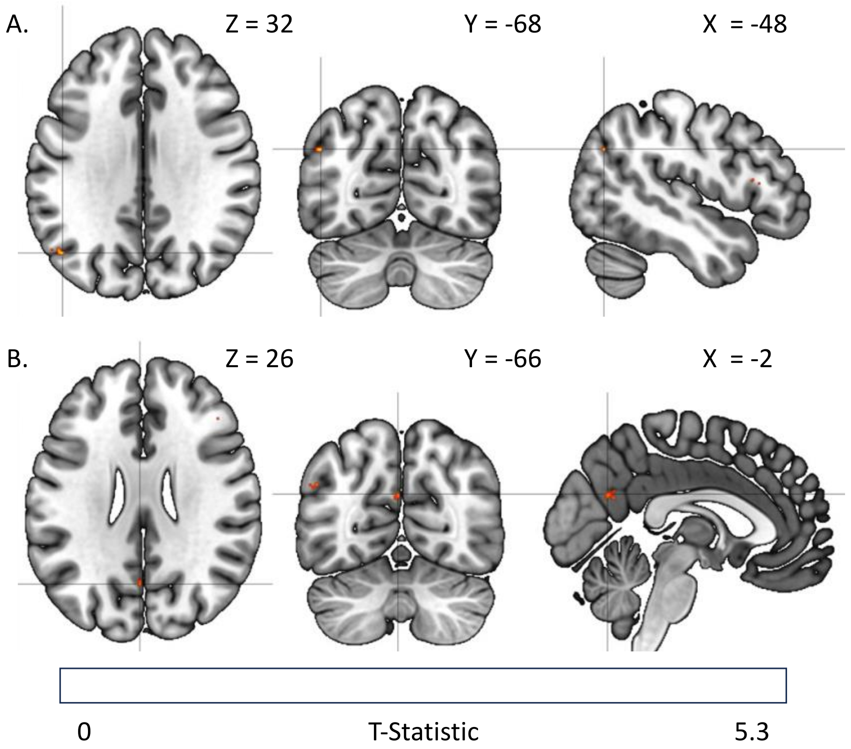


*Supplemental Figure 4.* Applying a liberal statistical threshold of $p < .01$, $K > 5$ voxels revealed clusters in the left angular gyrus (A) and left precuneus (B) associated with decoding visual perspective (i.e., in-body versus out-of-body) during memory retrieval. The results were rendered with 3D surface visualizations and projected onto individual slices onto the MNI152 T1-weighted template.

**Embodiment**. Although no regions survived *FWE*- correction when decoding embodiment during memory retrieval, a more liberal statistical threshold of $p < .01$, $K > 5$ voxels implicated the left angular gyrus and right precuneus, along with left posterior parietal regions (i.e., superior parietal gyrus and left intraparietal sulcus) and bilateral frontal regions (i.e., the right superior frontal gyrus corresponding to premotor cortex, bilateral precentral gyri, and bilateral middle frontal gyri) (see Supplemental Figure 5 and Supplemental Table 5).

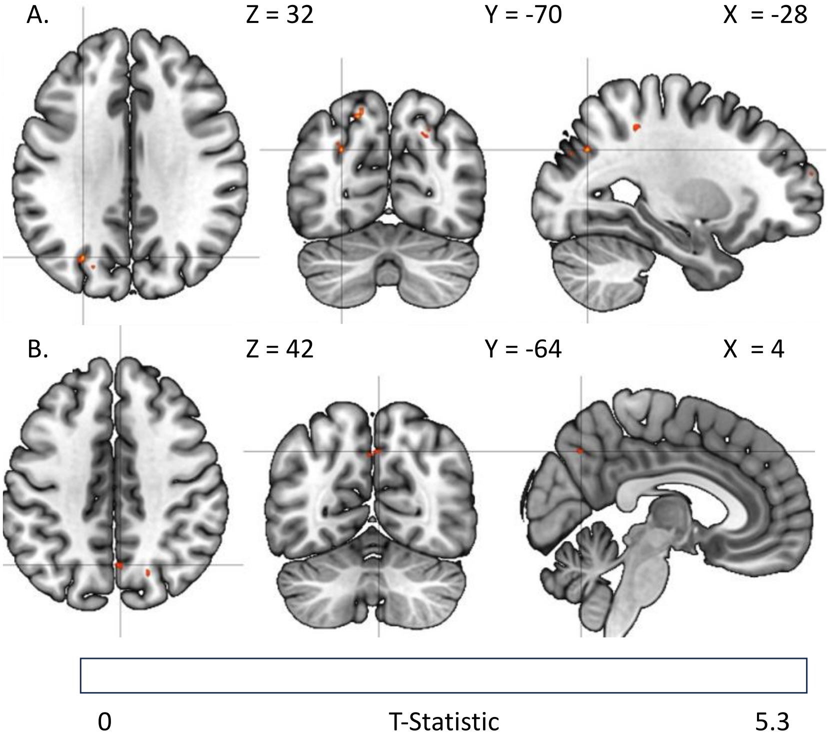


*Supplemental Figure 5.* Embodiment could be predicted from patterns of activity in the left angular gyrus (A) and right precuneus (B) at the statistical threshold of $p < .01$, $K > 5$ voxels. The results were rendered with 3D surface visualizations and projected onto individual slices onto the MNI152 T1-weighted template.

*Supplemental Table 5: Whole Brain Searchlight Analysis, Embodiment*

| Region | BA | K | $p_{uncorr}$ | T | X | Y | Z |
|---|---|---|---|---|---|---|---|
| Superior Parietal Gyrus | 7 | 7 | < .001 | 5.25 | -18 | -70 | 52 |
| Angular Gyrus | 39 | 5 | < .001 | 4.73 | -28 | -70 | 32 |
| Superior Frontal Gyrus | 10 | 7 | < .001 | 4.20 | -24 | 60 | 20 |
| Middle Temporal Gyrus | 21 | 14 | < .001 | 3.86 | -52 | -56 | 12 |
| Superior Occipital Gyrus | 7 | 10 | < .001 | 3.81 | 20 | -68 | 44 |
| Precuneus | 7 | 5 | .001 | 3.65 | 2 | -64 | 42 |
| Middle Occipital Gyrus | 7 | 13 | .001 | 3.64 | -22 | -76 | 30 |
| Middle Frontal Gyrus | 10 | 6 | .001 | 3.59 | -32 | 50 | 10 |
| Middle Frontal Gyrus | 10 | 5 | .001 | 3.35 | 42 | 50 | -6 |
| Intraparietal Sulcus | 7 | 5 | .002 | 3.28 | -28 | -42 | 46 |
| Corpus Callosum |  | 6 | .002 | 3.28 | -16 | 42 | 8 |
| Precentral Gyrus | 4 | 6 | .002 | 3.25 | -32 | -22 | 54 |
| Precentral Gyrus | 4 | 7 | .002 | 3.24 | 46 | -20 | 44 |
| Superior Frontal Gyrus | 6 | 5 | .002 | 3.18 | 8 | 30 | 64 |
| Superior Temporal Gyrus | 22 | 5 | .005 | 2.85 | 62 | -38 | 12 |

* The statistical map was thresholded at $p < .01$, $K > 5$ for display purposes. No clusters survived FWE-correction.

**Visual perspective x embodiment interaction**. Contrasting the synchronous in-body condition against the synchronous out-of-body condition did not identify any regions after correcting the statistical map at $p < .01$, $K > 5$ voxels. Among the asynchronous conditions, in-body versus out-of-body perspectives were linked to dissociable patterns of activity in the right supramarginal gyrus, left intraparietal sulcus, left inferior and middle frontal gyri, and right motor cortex at this liberal threshold (see Supplemental Figure 6 and Supplemental Table 6).

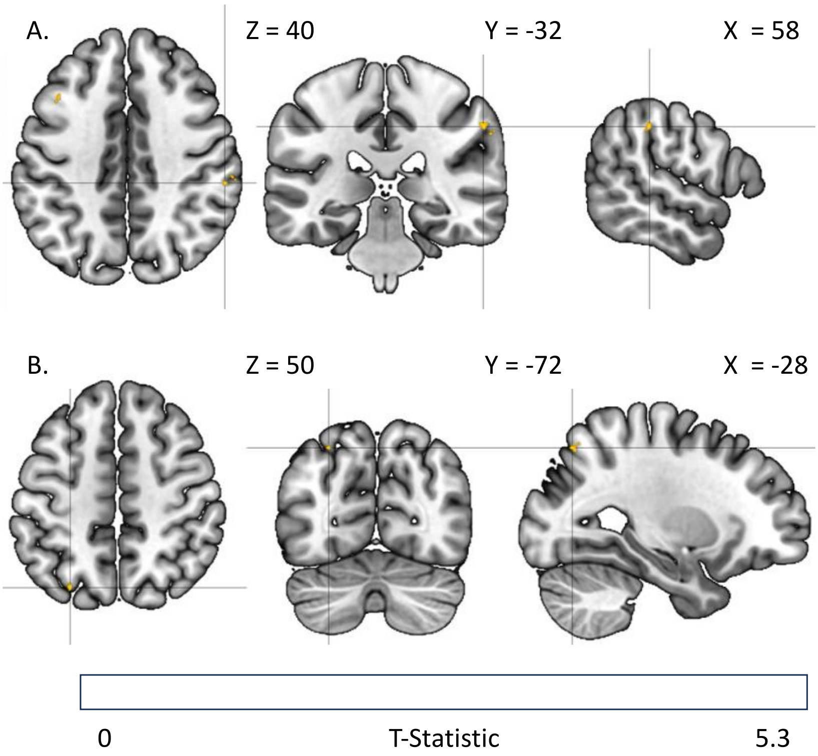


*Supplemental Figure 6.* Among asynchronous conditions, in-body compared to out-of-body perspectives were associated with dissociable patterns of activity in regions including the left supramarginal gyrus (A) and left intraparietal sulcus (B) at the threshold $p <. 01$, $K > 5$ voxels. The results were rendered with 3D surface visualizations and projected onto individual slices onto the MNI152 T1-weighted template.

| *Supplemental Table 6: Whole Brain Searchlight Analysis, Asynchronous In-Body vs. Asynchronous Out-of-Body* | | | | | | | |
|---|---|---|---|---|---|---|---|
| **Region** | **BA** | **K** | $p_{uncorr}$ | **T** | **X** | **Y** | **Z** |
| Supramarginal Gyrus | 40 | 5 | <.001 | 4.99 | 58 | -32 | 40 |
| Intraparietal Sulcus | 7 | 5 | <.001 | 4.42 | -28 | -72 | 50 |
| Supramarginal Gyrus | 40 | 7 | <.001 | 4.09 | 62 | -34 | 36 |
| Inferior Frontal Gyrus (Pars Triangularis) | 45 | 6 | .003 | 3.10 | -34 | 30 | 2 |
| Inferior Frontal Gyrus (Pars Orbitalis) | 47 | | .003 | 2.99 | -40 | 26 | -2 |
| Middle Frontal Gyrus | 8 | 56 | .003 | 2.96 | -40 | 18 | 40 |
| Precentral Gyrus | 4 | 5 | .004 | 2.89 | -38 | -18 | 58 |
| * The statistical map was thresholded at $p < .01$, $K > 5$ for display purposes. No clusters survived FWE-correction. | | | | | | | |
